# Targeting tau mitigates mitochondrial fragmentation and oxidative stress in amyotrophic lateral sclerosis

**DOI:** 10.1101/2021.03.22.436505

**Authors:** Tiziana Petrozziello, Evan A. Bordt, Alexandra N. Mills, Spencer E. Kim, Ellen Sapp, Benjamin A. Devlin, Abigail A. Obeng-Marnu, Sali M.K. Farhan, Ana C. Amaral, Simon Dujardin, Patrick M. Dooley, Christopher Henstridge, Derek H. Oakley, Andreas Neueder, Bradley T. Hyman, Tara L. Spires-Jones, Staci D. Bilbo, Khashayar Vakili, Merit E. Cudkowicz, James D. Berry, Marian DiFiglia, M. Catarina Silva, Stephen J. Haggarty, Ghazaleh Sadri-Vakili

## Abstract

Understanding the mechanisms underlying amyotrophic lateral sclerosis (ALS) is crucial for the development of new therapies. Recent evidence suggest that tau may be involved in ALS pathogenesis. Here, we demonstrated that hyperphosphorylated tau (pTau-S396) is mis-localized to synapses in human post-mortem motor cortex (mCTX) across ALS subtypes. Treatment with ALS synaptoneurosomes (SNs) derived from post-mortem mCTX, enriched in pTau-S396, increased oxidative stress, induced mitochondrial fragmentation, and altered mitochondrial connectivity *in vitro*. Furthermore, our findings revealed that pTau-S396 interacts with the pro-fission dynamin-related protein (DRP1), and similar to pTau-S396, DRP1 accumulated in ALS SNs across ALS subtypes. Lastly, reducing tau with a specific bifunctional degrader, QC-01-175, prevented ALS SNs-induced mitochondrial fragmentation and oxidative stress *in vitro*. Collectively, our findings suggest that increases in pTau-S396 may lead to mitochondrial fragmentation and oxidative stress in ALS and decreasing tau may provide a novel strategy to mitigate mitochondrial dysfunction in ALS.

**Graphical abstract:** 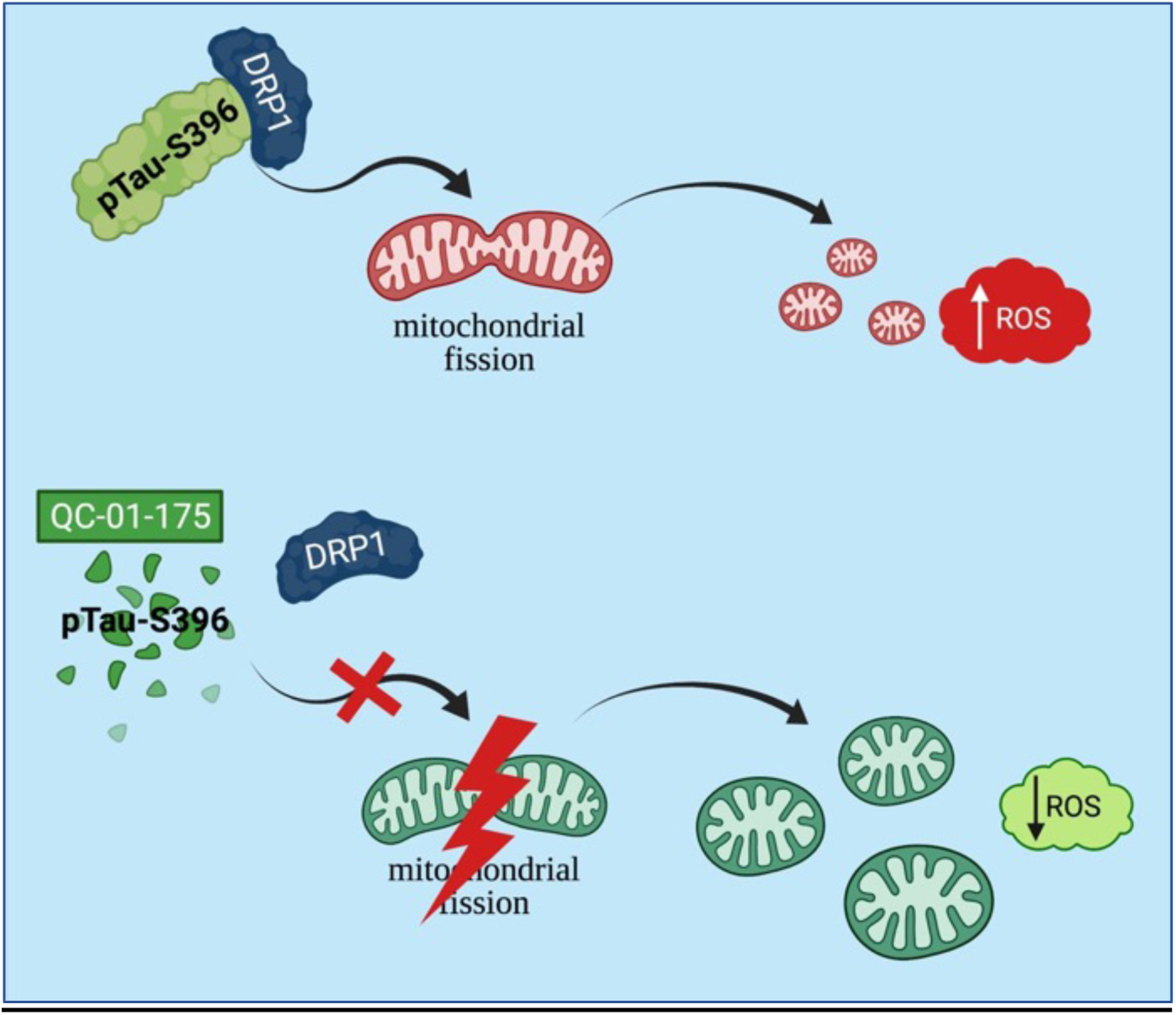

- pTau-S396 mis-localizes to synapses in ALS.
- ALS synaptoneurosomes (SNs), enriched in pTau-S396, increase oxidative stress and induce mitochondrial fragmentation *in vitro*.
- pTau-S396 interacts with the pro-fission GTPase DRP1 in ALS.
- Reducing tau with a specific degrader, QC-01-175, mitigates ALS SNs-induced mitochondrial fragmentation and increases in oxidative stress *in vitro*.

## Introduction

Amyotrophic lateral sclerosis (ALS) is a fatal neurodegenerative disease that primarily affects cortical and spinal motor neurons.^1^ Although 90% of ALS cases are sporadic, 10% are inherited and due to mutations in a number of genes such as superoxide dismutase 1 (*SOD1*), TAR DNA binding protein (*TARDBP*), fused in sarcoma (*FUS*), and a hexanucleotide repeat expansion in *C9ORF72* to name a few.^2^ Despite our understanding of the disease-causing mutations, the exact molecular mechanisms leading to motor neuron death remain unknown. Therefore, unraveling these mechanisms is crucial for the development of new therapeutic approaches.

Tau protein, a member of the microtubule-associated protein (MAP) family, plays a critical role in stabilizing microtubules, and is involved in several neuronal processes, including axonal transport, mobility and intracellular organization and trafficking of organelles.^3^ Recent studies have begun to link alteration in tau phosphorylation to ALS pathogenesis in both sporadic and familial cases,^4, 5^ as cytoplasmic inclusions of hyperphosphorylated tau have been described in post-mortem motor cortex (mCTX) and spinal cord from ALS patients.^6–8^

Tau is required for the trafficking of mitochondria across the axons to the synapses,^9^ a crucial event to sustain the high energy requirement of neuronal cells. Hyperphosphorylation of key epitopes on tau impairs this process and disrupts mitochondrial localization,^10–13^ thus contributing to axonal dysfunction and synapse loss in Alzheimer’s disease (AD).^9, 14–15^ Furthermore, tau over-expression and mis-localization decreases ATP levels, impairs oxidative phosphorylation (OXPHOS), and increases oxidative stress in AD and other neurodegenerative tauopathies.^16–18^ Lastly, accumulating evidence link hyperphosphorylated tau to impairment in mitochondrial dynamics,^17–18^ a physiological process that allows mitochondria to change their dynamic networks based on the energy requirement of the cell.^19^ Specifically, it has been reported that an abnormal interaction between hyperphosphorylated tau and dynamin-related protein 1 (DRP1), the GTPase involved in mitochondrial fission, exacerbates mitochondrial dysfunction in AD.^20–21^ Increases in oxidative stress and impairments in mitochondrial dynamics are also pathogenetic features and early events in the disease process in ALS.^1, 22–23^ Specifically, decreases in bioenergetics, calcium buffering, and mitochondrial transport as well as increases in oxidative stress have been described in animal and cellular models of ALS.^22–25^ Similarly, a decrease in cellular respiration, ATP production, and deficits in the mitochondrial electron transport chain have been described in post-mortem ALS spinal cord.^26–28^ Furthermore, mitochondria are structurally altered in ALS with the appearance of vacuolated and swollen mitochondria in ALS skeletal muscle and mCTX.^29–31^ Together these findings demonstrate that mitochondrial function and dynamics are altered in ALS; however, the exact mechanisms involved remain unknown. Here, we sought to determine whether tau contributes to alterations in mitochondrial morphology and functions in ALS.

## Results

### pTau-S396 is mis-localized to synapses in ALS independent of sex, region of onset, and genotype

Given that hyperphosphorylated tau accumulates at the synaptic level in AD,^32^ we first assessed the levels of tau and pTau-S396 in three different mCTX fractions – total, cytosolic, and synaptoneurosomes (SNs; the synaptic fraction) – derived from post-mortem ALS and controls by western blots. While the post-synaptic density protein 95 (PSD-95) was detected in the total and SNs fractions, it was absent in the cytosolic fraction derived from both ALS and control mCTX, demonstrating successful fractionation and purity of SNs. Furthermore, the results revealed that both tau and pTau-S396 mis-localized to SNs in ALS mCTX (Figure 1a). Specifically, while there was more tau in both control and ALS SNs compared to their cytosolic fractions (Figure 1b), pTau-S396 was only increased in ALS synapses (Figure 1c).

**Figure 1.**
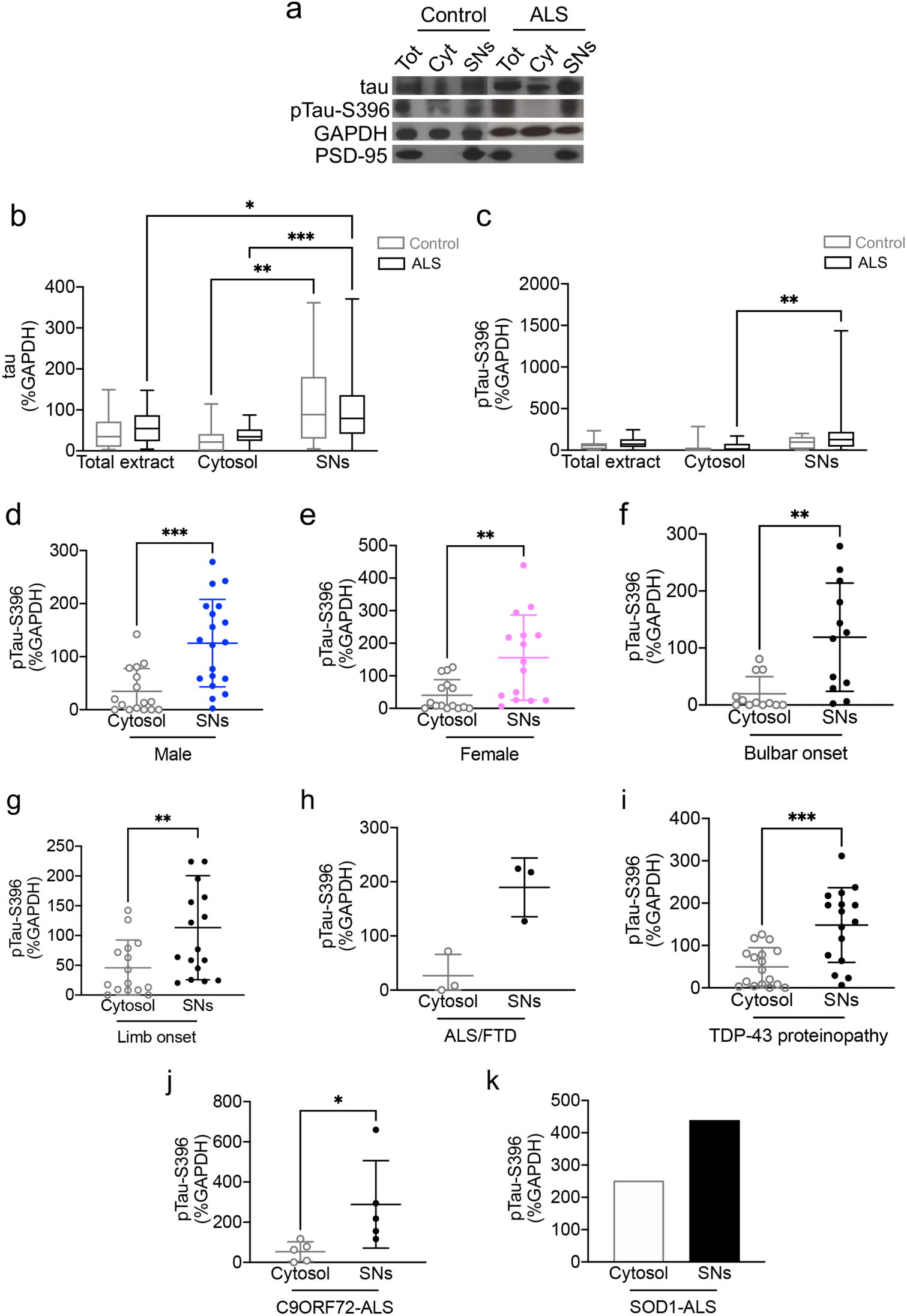
Synaptic pTau-S396 levels are increased independent of sex, region of onset, and genotype. (**a**) Representative western blot images of tau, pTau-S396 and PSD-95 in total, cytosolic and SNs fraction from ALS and control mCTX. (**b**) There was a significant effect of cellular fractions (two-way ANOVA, [F(2,137)=15.03], p<0.0001) on tau levels. Tau levels were significantly increased in control (n=14) and ALS SNs (n=36) compared to their cytosolic fractions (Tukey’s test, p=0.0059, and p=0.0006, respectively). There was a significant increase in tau levels in SNs compared to total extract in ALS (Tukey’s test, p=0.0489). (**c**) There was a significant effect of cellular fractions (two-way ANOVA, [F(2,133)=4.106], p=0.0186) on pTau-S396 levels together with a significant shift from ALS cytosolic fraction to the SNs (Tukey’s test, p=0.0024). There was a significant increase in synaptic pTau-S396 levels in (**d**) male (n=20; Mann-Whitney U test=49, p=0.0004) and (**e**) female ALS patients (n=16; Mann-Whitney U test=42, p=0.0026). Synaptic pTau-S396 levels were increased in (**f**) bulbar onset (n=13; Mann-Whitney U test=21, p=0.0023) and (**g**) limb onset ALS subjects (n=19; Mann-Whitney U test= 58, p=0.0077). (**h**) There was no change in pTau-S396 levels in ALS/FTD (n=3; Mann-Whitney U test=0, p=0.1000). (**i**) There was a significant increase in pTau-S396 in ALS with TDP-43 proteinopathy (n=18; Mann-Whitney U test=46, p=0.0008). (**j**) There was a significant increase in pTau-S396 in *C9ORF72*-ALS (n=5; Mann-Whitney U test=1, p=0.0159). (**k**) There was a significant increase in pTau-S396 in a single *SOD1*-ALS. *p<0.05; **p<0.01; ***p<0.001.

Next, we correlated increases in pTau-S396 in ALS SNs with the known patient clinicopathologic information. A significant shift in pTau-S396 from the cytosolic fraction to the SNs was observed in both male and female ALS mCTX (Figure 1d-e). Furthermore, synaptic pTau-S396 levels were significantly increased in both bulbar and limb onset ALS (Figure 1f-g). While there was no significant difference in synaptic pTau-S396 in ALS/frontotemporal dementia (ALS/FTD), there was a significant increase in SNs derived from ALS cases with TDP-43 proteinopathy (Figure 1h-i). There was a significant shift in pTau-S396 from the cytosol to SNs in *C9ORF72*-ALS, and the same shift was observed in the single *SOD1*-ALS case used in this study (Figure 1j-k).

### ALS SNs decrease cytosolic and mitochondrial aconitase activity without affecting cell survival

To verify whether ALS SNs, enriched in pTau-S396, may affect cell survival, human neuroblastoma SH-SY5Y cells were treated with SNs derived from control and ALS mCTX (10ng/mL) and recombinant tau protein as a positive control (10ng/mL) for 24h. Cell viability assessment by measuring active caspase-3 levels by an ELISA assay revealed no differences in the levels of active caspase-3 following treatment with either tau, control SNs, or ALS SNs (Figure 2a).

**Figure 2.**
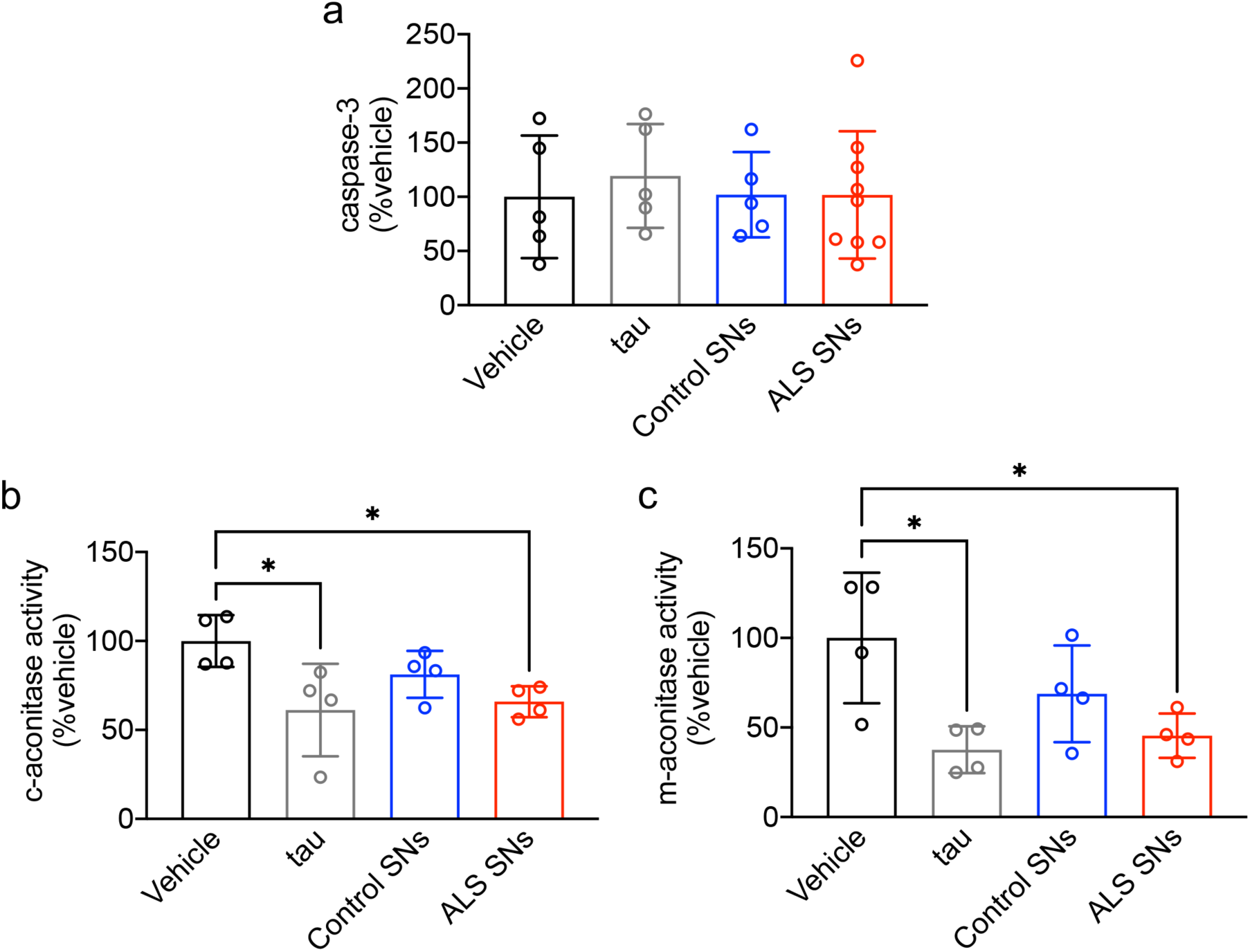
ALS SNs increase oxidative stress without affecting cell survival. (**a**) There was no effect of treatment on caspase-3 levels in SH-SY5Y cells (one-way ANOVA, [F(3,20)=0.1535], p=0.9262). Similarly, there were no change in caspase-3 levels in SH-SY5Y cells treated with tau (Tukey’s test, p=0.9203), control SNs (n=5; Tukey’s test, p=0.9999), or ALS SNs (n=9; Tukey’s test, p=0.999) compared to vehicle-treated cells. (**b**) There was a significant effect of treatment on c-aconitase activity in SH-SY5Y cells (one-way ANOVA, [F(3,12)=4.306], p=0.0280). c-aconitase activity was significantly decreased in tau- (Tukey’s test, p=0.0207) and ALS SNs-treated cells (n=4, Tukey’s test, p=0.0425) compared to vehicle-treated cells. (**c**) There was a significant effect of treatment on m-aconitase activity in SH-SY5Y cells (one-way ANOVA, [F(3,12)=5.269], p=0.0150). m-aconitase activity was significantly decreased in tau- (Tukey’s test, p=0.0106) and ALS SNs-treated cells (n=4, Tukey’s test, p=0.0245) compared to vehicle-treated cells. *p<0.05

Given that hyperphosphorylated tau has been associated with increased oxidative stress,^16–18^ we next assessed the activity of aconitase, a cellular enzyme that catalyzes the isomerization of citrate to isocitrate and is reversibly inactivated by increases in oxidative stress. We assessed the activity of both cytosolic and mitochondrial aconitase (c- and m-aconitase, respectively) in SH-SY5Y cells following treatment with tau, control and ALS SNs as well as antimycin A as positive control (25μM/30min) using an aconitase activity assay kit. As expected, antimycin A, an inhibitor of the mitochondrial complex III that increases ROS production,^33^ decreased both c- and m-aconitase activity (Suppl. Figure 1a-b). While there was no alteration following treatment with control SNs, there was a significant decrease in both c- and m-aconitase activity following treatment with recombinant tau or ALS-derived SNs compared to vehicle-treated cells (Figure 2b-c), indicative of increased oxidative stress.

### ALS SNs induce mitochondrial fragmentation

To determine whether ALS SNs may affect mitochondrial morphology, SH-SY5Y cells were treated with SNs derived from control and ALS mCTX (10ng/mL/24h) as well as recombinant tau protein (10ng/mL/24h) as positive control, and mitochondrial morphological parameters were measured by immunofluorescence using Tomm20 staining followed by volumetric reconstruction (Figure 3a). As expected, mitochondria within recombinant tau-treated cells had significantly decreased length and volume compared to mitochondria from vehicle- and control SNs-treated cells (Figure 3b-c). Similar decreases were observed in ALS SNs-treated cells compared to both vehicle- and control SNs-treated cells (Figure 3b-c). Specifically, there was a significant increase in the frequency of smaller mitochondria (<2 μm in length or <2 μm^3^ in volume) as well as a significant decrease in the frequency of larger mitochondria (>8μm in length or >10 μm^3^ in volume) in recombinant tau- or ALS SNs-treated cells compared to vehicle- or control SNs-treated cells (Suppl. Figure 2a-b).

**Figure 3.**
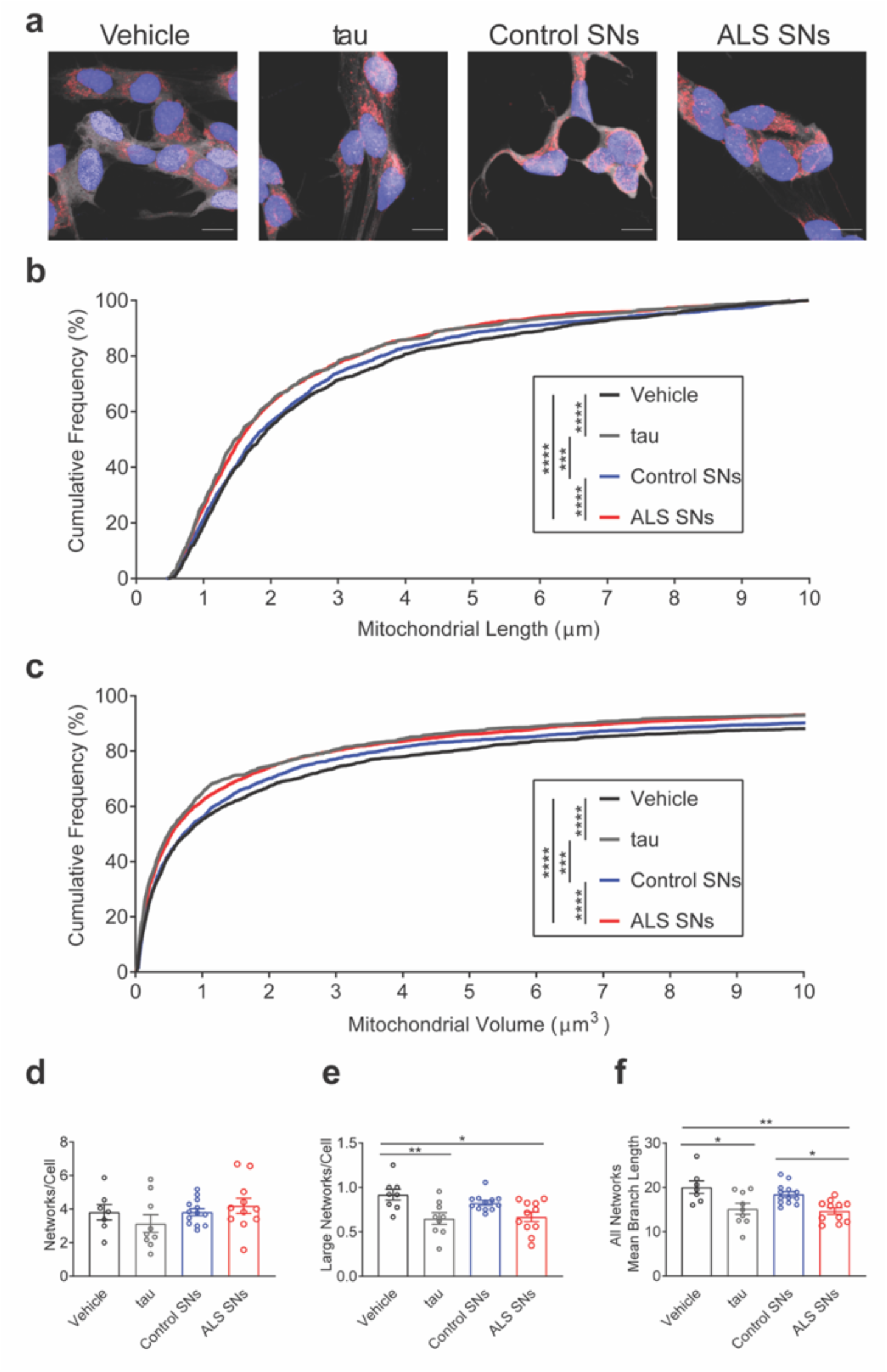
ALS SNs induce mitochondrial fragmentation. (**a**) Representative images of SH-SY5Y cells treated with vehicle, recombinant tau, control and ALS SNs stained with Hoechst (blue), CellMask (white), and Tomm20 (red). (**b**) Cumulative frequency graph indicated an effect of treatment on length (Kruskal-Wallis, H=55.78, p<0.0001) with smaller mitochondria in recombinant tau- and ALS SNs- (n=3) compared to vehicle- (p<0.0001, and p<0.0001, respectively) and control SNs (n=3)-treated cells (p=0.0001, and p<0.0001, respectively). (**c**) Cumulative frequency graph demonstrated an effect of treatment on volume (Kruskal-Wallis, H=51.2941, p<0.0001) with smaller mitochondria in recombinant tau- and ALS SNs-compared to vehicle- (p<0.0001, and p<0.0001, respectively) and control SNs-treated cells (p=0.0001, and p<0.0001, respectively). (**d**) Treatment with either recombinant tau or ALS SNs had no effect on mitochondrial networks/cell (one-way ANOVA, [F(3,36)=1.157], p=0.3396). (**e**) There was an effect of treatment on large networks/cell (one-way ANOVA, [F(3,36)=5.809], p=0.0024). Tukey’s test demonstrated a decrease in recombinant tau- or ALS SNs-compared to vehicle-treated cells (p=0.0082, and p=0.0105, respectively). (**f**) There was an effect of treatment on the mean branch length of mitochondria within networks (one-way ANOVA, [F(3,36)=6.901], p=0.0009). Tukey’s test revealed a decrease in recombinant tau-compared to vehicle-treated cells (p=0.0124) and in ALS SNs-compared to vehicle- and control SNs-treated cells (p=0.0032, and p=0.0184, respectively). *p<0.05; **p<0.01; ***p<0.001, ****p<0.0001

In addition to morphometric analyses of individual mitochondrial morphologies, we also examined the ability of ALS SNs to affect connectivity of the mitochondrial networks. While treatment with either tau or ALS SNs had no effect on the number of mitochondrial networks/cell (Figure 3d), there was a significant decrease in large networks/cell in recombinant tau- or ALS SNs-treated cells compared to vehicle-treated cells (Figure 3e). Consistent with individual mitochondrial analyses, there was also a significant decrease in the mean mitochondrial branch length in recombinant tau-treated cells compared to vehicle-treated cells as well as in ALS SNs-treated cells compared to both vehicle- and control SNs-treated cells (Figure 3f).

### DRP1 interacts with pTau-S396 and its levels are increased in ALS SNs

Given that pTau-S396 has been shown to interact with the pro-mitochondrial fission GTPase DRP1,^20^ and ALS SNs induce mitochondrial fragmentation *in vitro*, we hypothesized that increases in pTau-S396 may trigger pathological mitochondrial fission in ALS by binding DRP1. To test this hypothesis, we first assessed the possible interaction between pTau-S396 and DRP1 in control and ALS post-mortem mCTX by co-immunoprecipitation. Our results revealed that pTau-S396 interacts with DRP1 in mCTX, however there was no significant difference in this interaction between control and ALS (Figure 4a-b).

**Figure 4.**
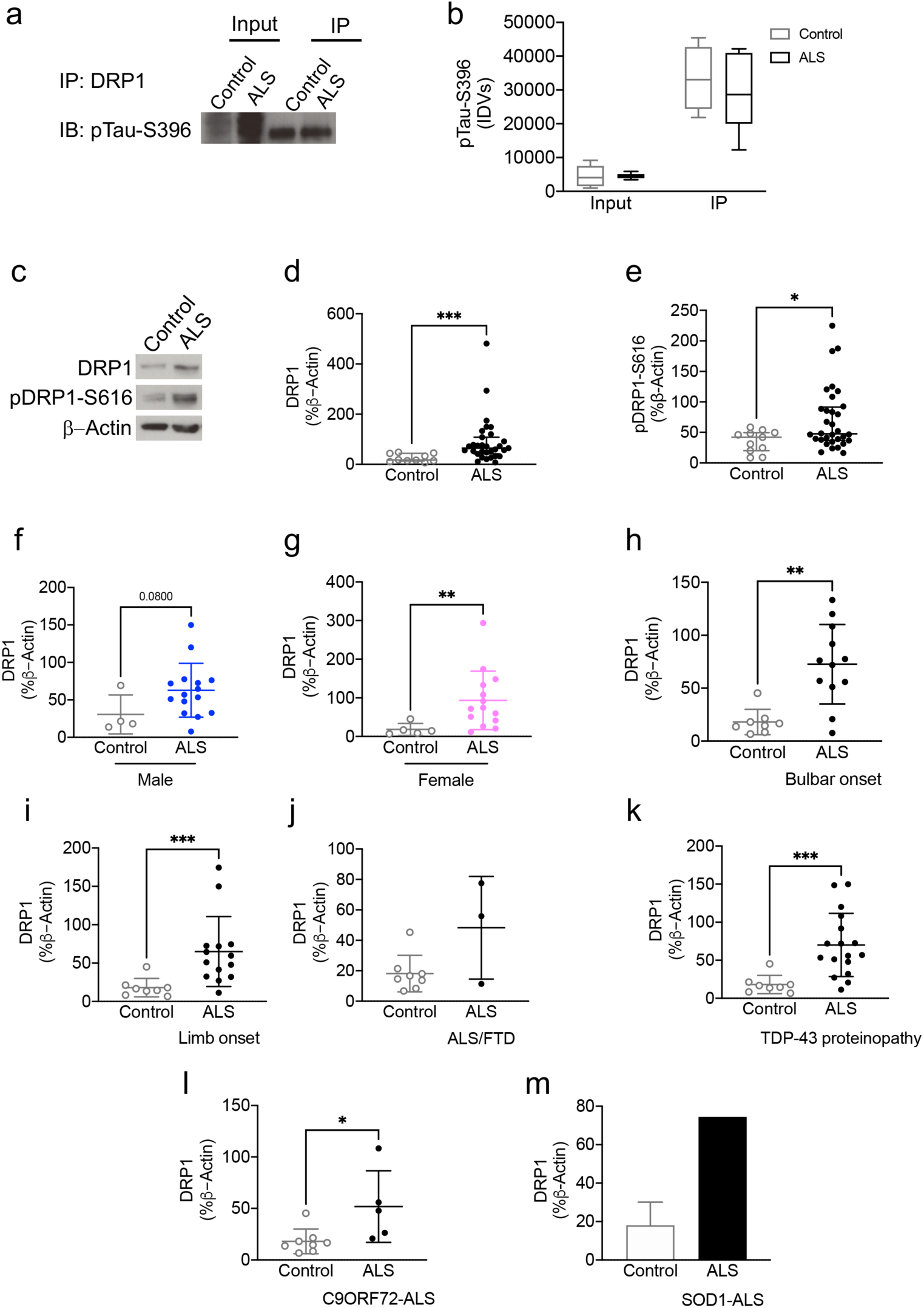
DRP1 interacts with pTau-S396 and its levels are increased in ALS SNs. **(a)** Representative western blot images of pTau-S396 and DRP1 interaction in control and ALS mCTX. **(b)** There was a significant effect of cellular fractions in pTau-S396 and DRP1 interaction (two-way ANOVA, [F(1,27)=92.60], p<0.0001) with no significant changes in this interaction between control (n=8) and ALS mCTX (n=8) (Tukey’s test, p=0.6958). (**c**) Representative western blot images of DRP1 and pDRP1-S616 in ALS and control SNs. There was a significant increase in (**d**) DRP1 (Mann-Whitney U test=41, p=0.0002) and (**e**) pDRP1-S616 levels (Mann-Whitney U test=103.5, p=0.0429) in ALS SNs (n=32) compared to controls (n=12). (**f**) There was a trend towards a significant increase in synaptic DRP1 levels in male (n=16) ALS patients compared to controls (n=3) (Mann-Whitney U test=12, p=0.0800). (**g**) Synaptic DRP1 levels were increased in female ALS patients (n=15) compared to controls (n=5) (Mann-Whitney U test=6, p=0.0050). Synaptic DRP1 levels were increased in (**h**) bulbar onset (n=13; Mann-Whitney U test=9, p=0.0015) and (**i**) limb onset ALS (n=15; Mann-Whitney U test=10, p=0.0009) compared to controls (n=9). (**j**) There was no change in DRP1 levels in ALS/FTD (n=3; Mann-Whitney U test=6, p=0.2788). (**k**) There was a significant increase in DRP1 in ALS with TDP-43 proteinopathy (n=18; Mann-Whitney U test=10, p=00002). (**l**) There was a significant increase in pTau-S396 in *C9ORF72*-ALS (n=5; Mann-Whitney U test=3, p=0.0109). (**m**) There was a significant increase in pTau-S396 in a single *SOD1*-ALS. *p<0.05; **p<0.01; ***p<0.001.

Given the interaction between pTau-S396 and DRP1 and the increase in synaptic pTau-S396 levels in ALS, we next assessed the levels of mitochondrial fission and fusion markers in SNs derived from ALS and control mCTX by western blots. The results revealed a significant increase in both DRP1 and its active form pDRP1-S616 in ALS SNs compared to control SNs (Figure 4c-e), while there was no significant difference in the levels of pro-fusion proteins, Mfn1, Mfn2, or OPA1 (Suppl. Figure 3a-d), suggesting an increase in mitochondrial fission in ALS.

As pTau-S396 mis-localization at the synaptic level occurs across subtypes of ALS, we correlated increases in DRP1 in ALS SNs with the known patient clinicopathologic information. The results revealed a trend towards a significant increase in DRP1 levels in SNs-derived from male ALS patients as well as a significant increase in female patients (Figure 4f-g). Furthermore, synaptic DRP1 levels were significantly increased in both bulbar and limb onset ALS (Figure 4h-i). While there were no alterations in synaptic DRP1 levels in ALS/FTD, there was a significant increase in SNs from ALS cases with TDP-43 proteinopathy (Figure 4j-k). Lastly, there was a significant increase in synaptic DRP1 levels in *C9ORF72*-ALS, and the same increase was observed in the single *SOD1*-ALS subject used in this study (Figure 4l-m).

### Mitochondrial morphology is altered in ALS post-mortem mCTX

Given the significant increase in DRP1, we next assessed the levels of the mitochondrial outer membrane marker Tomm20 by western blots in post-mortem ALS and control mCTX. The data revealed a significant decrease in Tomm20 levels in ALS compared to control mCTX, suggesting alterations in mitochondrial density, mass, or morphology (Figure 5a). Similarly, analysis of post-mortem mCTX using electron microscopy revealed a significant decrease in the density as well as in the length of mitochondria (Figure 5b, white arrow) in ALS compared to controls (Figure 5c-d). Furthermore, there was a significant increase in the frequency of shorter mitochondria (<0.50μm) in ALS mCTX (Figure 5e), consistent with increased mitochondrial fragmentation in ALS.

**Figure 5.**
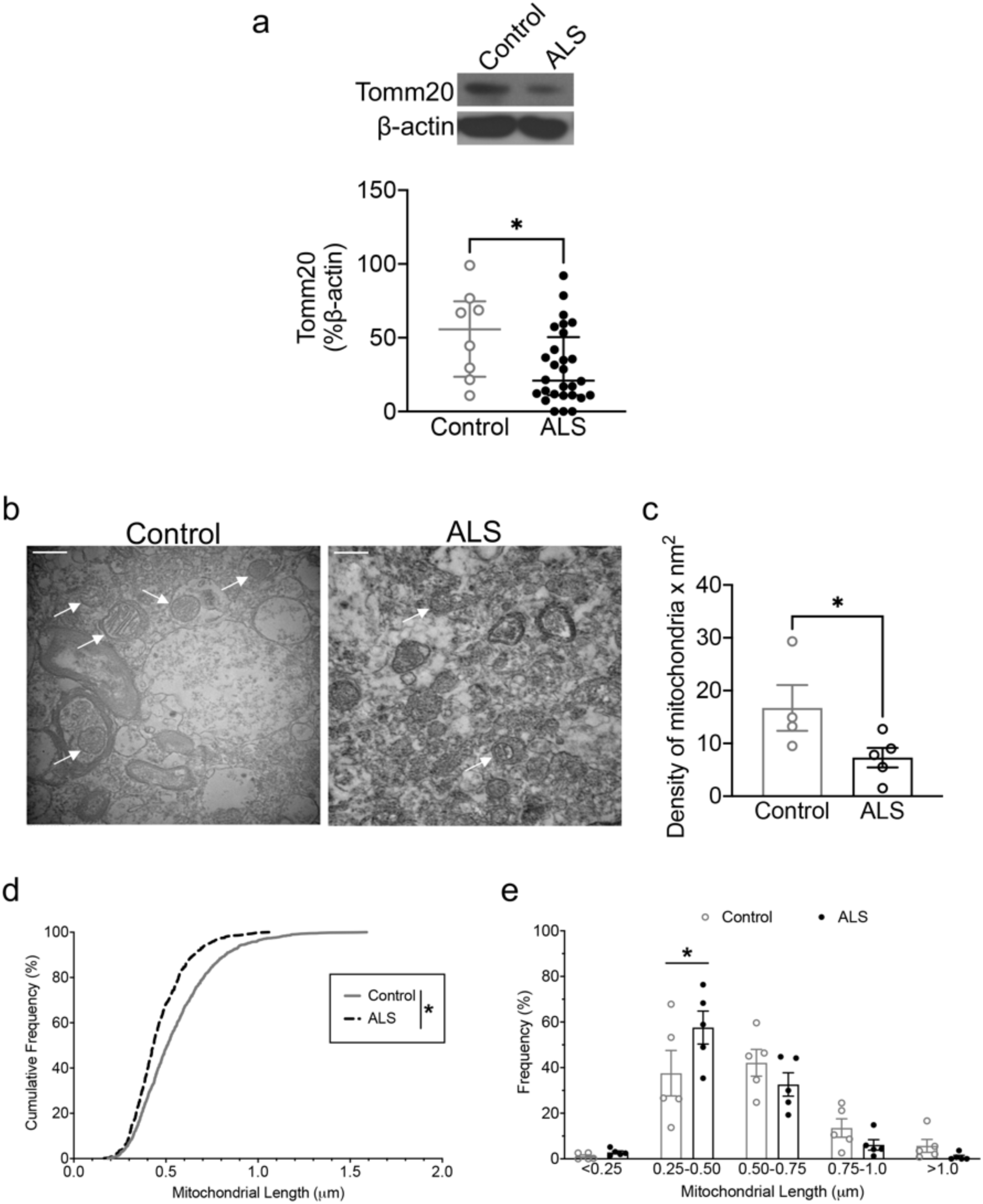
Mitochondrial mass, density and length are decreased in ALS mCTX. (**a**) There was a significant decrease in Tomm20 levels in ALS mCTX (n=30) compared to controls (n=10) (Mann-Whitney U test=60.50, p=0.0495). (**b**) Representative EM images from control and ALS mCTX. There were fewer and smaller mitochondria (white arrows) in ALS compared to control mCTX. (**c**) There was a significant decrease in the density of mitochondria in ALS mCTX (n=5) compared to controls (n=4) (Mann-Whitney U test=1, p=0.0317). (**d**) There was a significant decrease in the length of mitochondria in ALS mCTX compared to controls (Kolmogorov-Smirnov test, p<0.0001). (**e**) There was a significant effect of mitochondrial length (two-way ANOVA, [F(3,32)=30.08], p<0.0001) and length X disease interaction (two-way ANOVA, [F(3,32)=3.314], p=0.0032) in mCTX. Furthermore, there was a significant increase in the frequency of smaller mitochondria (<0.50um) in ALS mCTX compared to controls, while there was no change in the frequency of larger mitochondria (Tukey’s test, p=0.0432 and p>0.9999, respectively). Scale bar: 500nm. *p<0.05.

### siDRP1 mitigates ALS SNs-induced alterations in mitochondrial morphology

Given that DRP1 inhibition prevents mitochondrial dysfunction in AD,^21^ we used a siRNA approach to reduce *DRP1* levels in SH-SY5Y cells prior to treatment with control or ALS SNs. SH-SY5Y cells were treated with 10, 100 or 150nM of siDRP1 for 24 and 48h to identify the effective dose and treatment time for knocking down *DRP1*. Our results demonstrated a significant decrease in DRP1 levels following silencing, with no signal detected following exposure to 150nM siDRP1 for 24h or following 48h treatment (Suppl. Figure 4a). Therefore, SH-SY5Y cells were transfected with 150nM of siDRP1 for 24h and then treated with ALS SNs (10ng/mL/24h) before assessing DRP1 levels by western blots. The results demonstrated a significant decrease in DRP1 levels in both siDRP1- and siDRP1+ALS SNs-treated cells compared to both siControl- and siControl+ALS SNs-treated cells (Suppl. Figure 4b).

For mitochondrial morphological assessments, SH-SY5Y cells were silenced with siDRP1 (150nM/24h) followed by treatment with recombinant tau or SNs derived from control and ALS mCTX (10ng/mL/24h) (Suppl. Figure 5a). As expected, treatment with both recombinant tau and ALS SNs in the presence of siControl significantly reduced mitochondrial length and volume, as mitochondria were significantly shorter in recombinant tau- and ALS SNs-treated cells compared to siControl- or siControl+control SNs- treated cells (Suppl. Figure 5b-c). Furthermore, DRP1 knockdown prevented recombinant tau- and ALS SNs-induced increases in smaller mitochondria and reductions in larger mitochondria. Specifically, the treatment with ALS SNs resulted in fewer smaller mitochondria and more larger mitochondria in siDRP1-compared to siControl-treated cells (Suppl. Figure 5b-c). Similarly, silencing DRP1 prevented the increase in the frequency of smaller mitochondria (<2μm in length and <2μm^3^ in volume) as well as the decrease in the frequency of larger mitochondria (>8μm in length and >10 μm^3^in volume) induced by recombinant tau or ALS SNs (Suppl. Figure 6a-b).

Mitochondrial connectivity analyses led to similar observations, with no differences in the number of overall networks/cell (Suppl. Figure 5d). Similar to length and volume measurements, knocking down *DRP1* prevented recombinant tau- and ALS SNs-induced alterations in large networks/cell, as there was a significant increase in the number of large networks/cell in siDRP1-exposed cells treated with either recombinant tau or ALS SNs (Suppl. Figure 5e). Lastly, siDRP1 prevented ALS SNs-mediated decrease in branch length, as siDRP1+ALS SNs-treated cells had longer branch length compared to siControl+ALS SNs-treated cells (Suppl. Figure 5f).

### Tau degradation mitigates ALS SNs-induced alterations in mitochondrial morphology

To test the hypothesis that reducing tau may recapitulate the same effect induced by knocking down *DRP1* on mitochondrial morphology, a novel bifunctional tau degrader, QC-01-175, which has been shown to be capable of recruiting tau to the E3 ubiquitin ligase CRL4^CRBN^ resulting in its degradation from the proteasome,^34^ was used to reduce pTau-S396 levels in SH-SY5Y cells following treatment with control or ALS SNs. To identify the effective dose and treatment time for QC-01-175 able to reduce pTau-S396 levels, SH-SY5Y cells were first treated with 1 or 10μM of QC-01-175 for 2, 4, and 24h following treatment with ALS SNs (10ng/mL/24h). Our results revealed that while 1μM QC-01-175 had no effect on pTau-S396 levels, 10μM of the degrader reduced pTau-S396 in ALS SNs-treated cells compared to vehicle-treated cells following 4h and that this effect was absent following 24h at the same concentration (Figure 6a-b). Therefore, SH-SY5Y cells were treated with ALS SNs (10ng/mL/24h) followed by QC-01-175 (10μM) for 4h before assessing pTau-S396 levels by western blots. The results revealed a significant decrease in pTau-S396 levels in cells treated with QC-01-175 following exposure to ALS SNs compared to ALS SNs-treated cells (Figure 6c).

**Figure 6.**
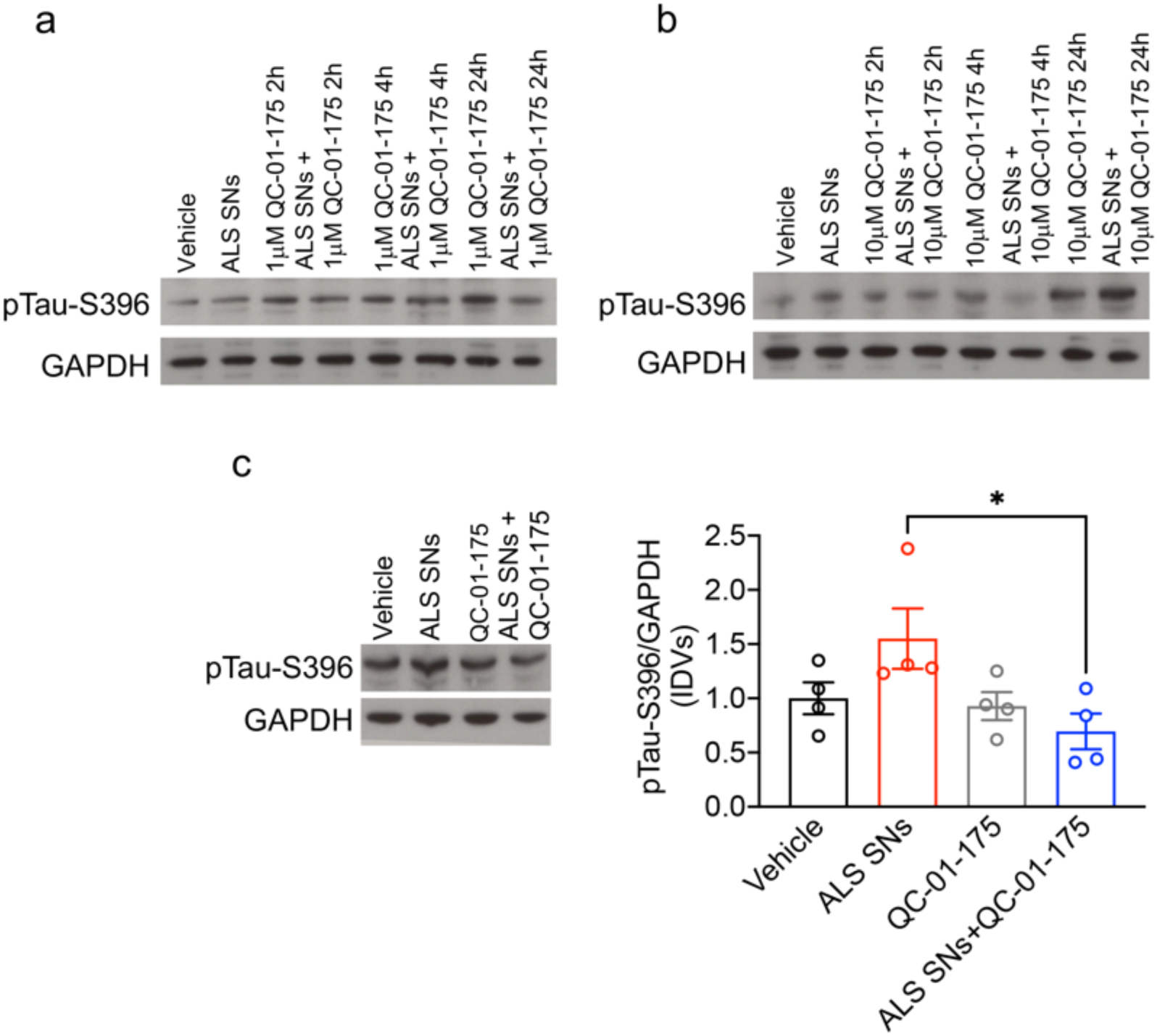
QC-01-175 decreases pTau-S396 levels in ALS SNs-treated cells. (**a**) Representative western blot images of pTau-S396 in vehicle- and ALS SNs-treated SH-SY5Y cells following treatment with QC-01-175 (1μM) for 2, 4 or 24h. (**b**) Representative western blot images of pTau-S396 in vehicle- and ALS SNs-treated SH-SY5Y cells treated with QC-01-175 (10μM) for 2, 4 or 24h. pTau-S396 levels were decreased following treatment with QC-01-175 (10μM) for 24h. (**c**) There was a significant effect of treatment on pTau-S396 levels in SH-SY5Y cells (one-way ANOVA, [F(3,12)=3.699], p=0.0429) with a significant decrease in pTau-S396 levels in ALS SNs+QC-01-175-treated cells compared to ALS SNs-treated cells (Tukey’s test, p=0.0328). *p<0.05.

For mitochondrial morphological assessments, SH-SY5Y cells were treated with either recombinant tau or SNs derived from control and ALS mCTX (10ng/mL/24h) followed by treatment with QC-01-175 (10μM/4h) before assessing mitochondrial morphological parameters (Figure 7a). As in previous experiments, recombinant tau and ALS SNs treatment significantly shortened mitochondrial length and volume compared to vehicle- and control SNs-treated cells. Degrading tau with QC-01-175 reverted recombinant tau and ALS SNs-induced increases in smaller mitochondria and decreases in larger mitochondria. Specifically, there were fewer smaller mitochondria and more larger mitochondria in QC-01-175+ALS SNs-treated cells compared to ALS SNs-treated cells (Figure 7b-c). Furthermore, selective degradation of tau using QC-01-175 prevented recombinant tau- and ALS SNs-induced mitochondrial length and volume alterations, as treatment with tau or ALS SNs resulted in fewer smaller mitochondria (<2μm in length and <2μm^3^ in volume) following QC-01-175 treatment (Suppl. Figure 7a-b). Lastly, there were more larger mitochondria (>8μm^3^ in volume) in ALS SNs+QC-01-175-treated cells compared to ALS SNs-treated cells (Suppl. Figure 7b).

**Figure 7.**
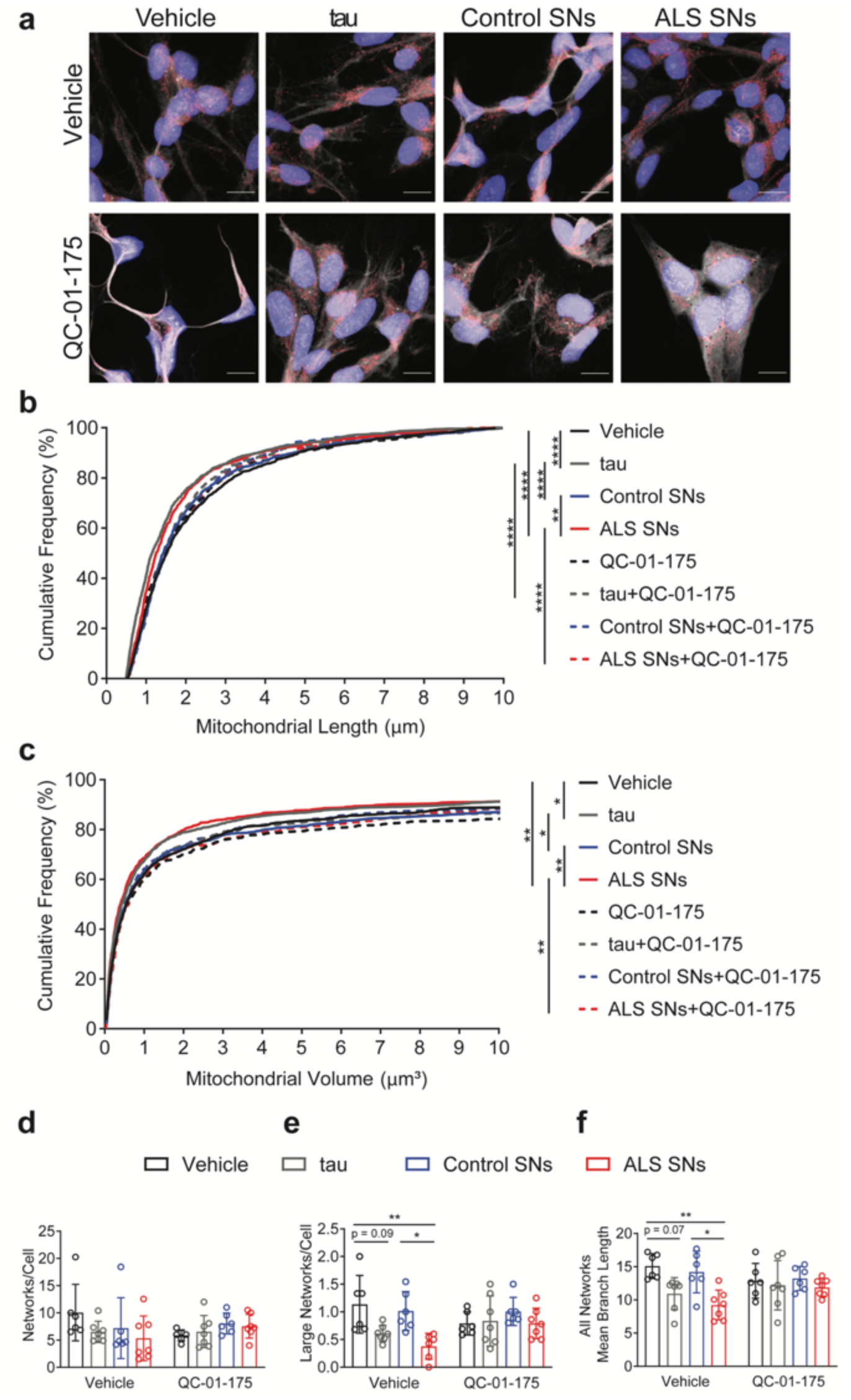
Degrading tau mitigates ALS SNs-induced mitochondrial fragmentation. (**a**) Representative image of SH-SY5Y cells treated with vehicle, recombinant tau, control and ALS SNs in the absence or presence of QC-01-175 stained with Hoechst (blue), CellMask (white), and Tomm20 (red). (**b**) Cumulative frequency graph indicated an effect of treatment on length (Kruskal-Wallis, H=113.3, p<0.0001) with smaller mitochondria in tau- and ALS SNs- (n=3) compared to vehicle- (p<0.0001, and p<0.0001, respectively) and control SNs (n=3)-treated cells (p<0.0001, and p=0.0015, respectively). QC-01-175 prevented decrease in mitochondrial length induced by both tau (p<0.0001) and ALS SNs (p=0.0170). (**c**) Cumulative frequency graph indicated an effect of treatment on volume (Kruskal-Wallis, H=39.11, p<0.0001) with smaller mitochondria in tau- and ALS SNs-compared to vehicle- (p=0.0081, and p=0.0039, respectively) and control SNs-treated cells (p=0.0112, and p=0.0024, respectively). QC-01-175 prevented decrease in mitochondrial volume induced by ALS SNs (p=0.0020). (**d**) There was no effect of treatment on total networks/cell following treatment with tau, control and ALS SNs (two-way ANOVA, [F(1,44)=0.07953], p=0.7793). (**e**) There was an effect of treatment (two-way ANOVA, [F(3,43)=4.792] p=0.0057) and treatment X tau degradation (two-way ANOVA [F(3,43)=3.121], p=0.0357) in SH-SY5Y cells with a significant decrease in ALS SNs-compared to vehicle- or control SNs-treated cells (Tukey’s test, p=0.0052, and p=0.0307, respectively). (**f**) There was an effect of treatment (two-way ANOVA, [F(3,44)=5.770], p=0.0020) on mean mitochondrial branch length within networks with a decrease in ALS SNs-compared to vehicle- or control SNs-treated cells (Tukey’s test, p=0.0028, and p=0.0182, respectively). *p<0.05; **p<0.01; ****p<0.0001.

Mitochondrial network analyses revealed no alterations in the number of total networks/cell, similar to previous experiments (Figure 7d). There were also no alterations in the number of large networks or mitochondrial branch length following treatment with QC-01-175 (Figure 7e-f). Although QC-01-175 treatment more than doubled the number of large networks compared to ALS SNs treatment, this protection did not rise to statistical significance.

### Tau degradation prevents the increase in reactive oxygen species induced by ALS SNs

Given that treatment with ALS SNs increases oxidative stress, and QC-01-175 ameliorates mitochondrial morphology, we next verified whether QC-01-175 was also able to mitigate oxidative stress induced by ALS SNs. Therefore, SH-SY5Y cells were treated with either recombinant tau or SNs derived from control and ALS mCTX (10ng/mL/24h) followed by treatment with QC-01-175 (10μM/4h) before assessing reactive oxygen species (ROS) levels by using CellROX, a fluorogenic probe for measuring oxidative stress in live cells (Figure 8a). Antimycin A (25μM/30min) was used as positive control (Suppl. Figure 8). According to previous results, the treatment with both recombinant tau and ALS SNs induced an increase in ROS levels compared to both vehicle- and control SNs-treated cells (Figure 8b). Importantly, ROS levels were significantly decreased in both recombinant tau+QC-01-175- and ALS SNs+QC-01-175-treated cells compared to recombinant tau- and ALS SNs-treated cells (Figure 8b), indicating a rescue of oxidative stress by tau-specific clearance.

**Figure 8.**
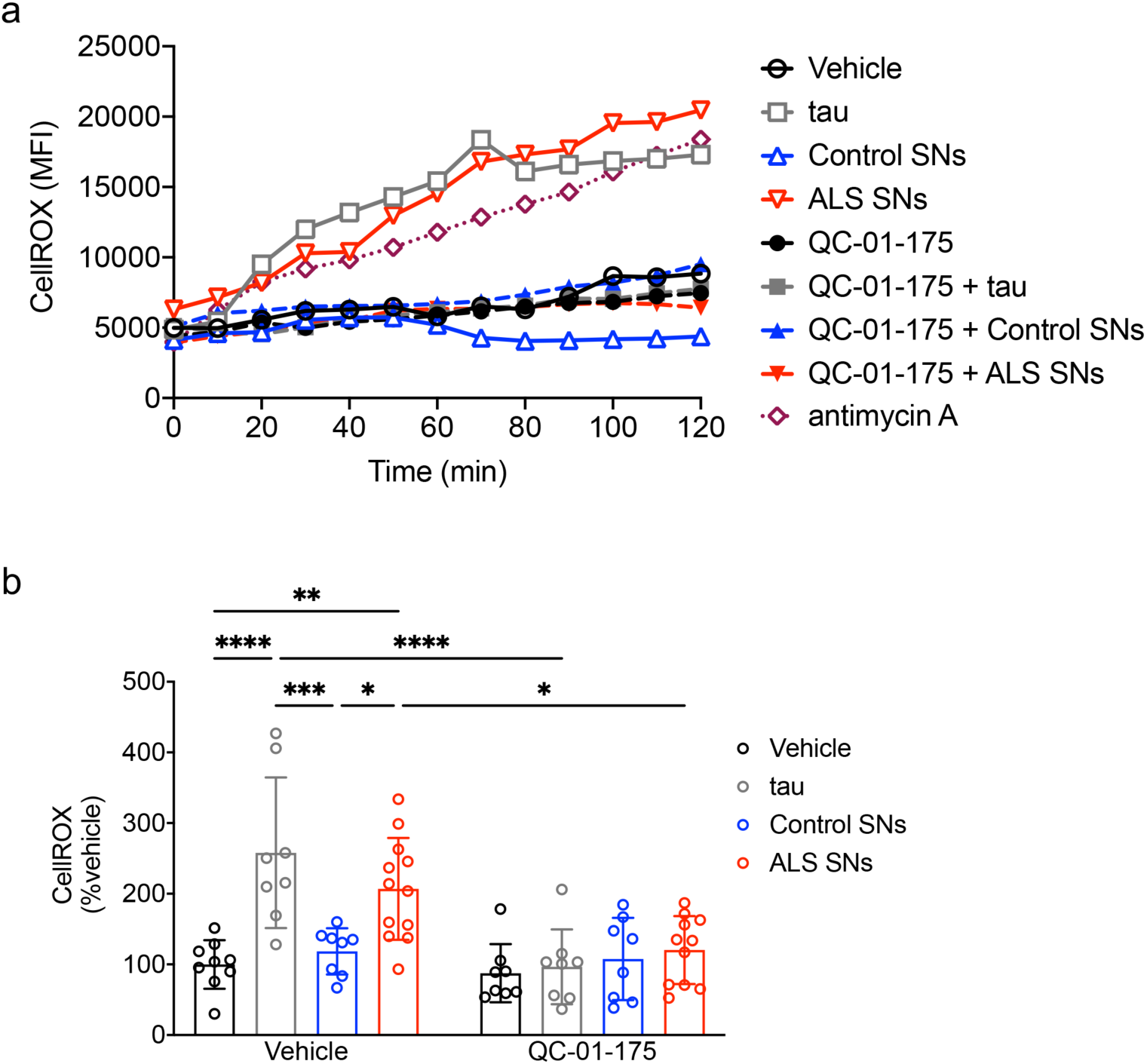
QC-01-175 prevents ALS SNs-induced increases in ROS levels. (**a**) Representative traces of CellRox mean fluorescence intensity (MFI) measured over time in SH-SY5Y cells treated with recombinant tau, control and ALS SNs (10ng/mL/24h) followed by exposure to QC-01-175 (10μM/4h) and incubated with CellRox (2.5μM/30min). (**b**) There was a significant effect of treatment (two-way ANOVA, [F(3,64)=7.616], p=0.0002), QC-01-175 (two-way ANOVA, [F(3,64)=22.27], p<0.0001), and treatment X QC-01-175 (two-way ANOVA, [F(3,64)=5.801], p=0.0014) on ROS levels. There was an increase in ROS levels in recombinant tau- and ALS SNs- (n=9) compared to vehicle-(Tukey’s test, p<0.0001, and p=0.0036, respectively) and control SNs (n=4)-treated cells (Tukey’s test, p=0.0005, and p=0.0398, respectively). QC-01-175 prevented increase in ROS levels induced by recombinant tau and ALS SNs (Tukey’s test, p<0.0001, and p=0.0213, respectively). *p<0.05; **p<0.01; ***p<0.001; ****p<0.0001.

## Discussion

In this study we reported for the first time mis-localization of hyperphosphorylated tau at S396 to synapses across subtypes of ALS in a large cohort of human post-mortem ALS mCTX. We also demonstrated that ALS SNs, enriched in pTau-S396, increase oxidative stress *in vitro* without affecting cell survival. Treatment with ALS SNs was also sufficient to alter mitochondrial length, volume, and connectivity *in vitro*, thus indicating increases in mitochondrial fragmentation. Although pTau-S396 was found to interact with DRP1 in both control and ALS mCTX, there was only a significant increase in DRP1 levels in ALS SNs that was observed across subtypes similar to pTau-S396. Our results also revealed a decrease in mitochondrial density and length in ALS mCTX, supporting previous findings.^22-23^ Importantly, we demonstrated that reducing tau with the specific tau degrader, QC-01-175,^34^ mitigated alterations in mitochondrial morphology, similar to knocking down *DRP1*, and decreased ROS levels, thus suggesting a role for tau in inducing mitochondrial fragmentation and oxidative stress in ALS.

Recently, alterations in tau phosphorylation have been reported in both sporadic and familial cases of ALS.^4–5^ Specifically, cytoplasmic inclusions of hyperphosphorylated tau have been described in ALS post-mortem mCTX and spinal cord.^6–8^ Here, our results expand on these findings and demonstrate a significant mis-localization of tau and pTau-S396 from the cytosol to synapses, reminiscent of AD.^32^ Furthermore, this synaptic mis-localization occurred across ALS subtypes, suggesting that hyperphosphorylation of tau may represent a common mechanism underlying ALS pathogenesis; however, further investigation is required to determine the pathology of genetic subtypes with a larger sample size. Additional studies assessing tau phosphorylation at different epitopes are also necessary to better characterize the involvement of tau hyperphosphorylation, mis-localization and toxicity in ALS pathogenesis.

Although the exact molecular mechanisms underlying tau toxicity are not yet fully understood, several studies suggest that tau may trigger mitochondrial dysfunction in neurodegenerative diseases, such as AD.^17–18^ Mitochondrial loss and dysfunction lead to tau hyperphosphorylation and aggregation, suggesting a deleterious interplay between tau and mitochondria that may contribute to alterations in neuronal and synaptic function in neurodegeneration.^17^ Specifically, tau hyperphosphorylation has been shown to alter mitochondrial localization at the neuronal level,^10–13, 15^ and, in turn, axonal mitochondrial loss further exacerbates tau hyperphosphorylation and its disassembly from the microtubules within the axons.^17, 35^ Mitochondria are mis-localized in both animal and cellular models of ALS carrying mutations in *SOD1* and *TARDBP*,^36–40^ and have been shown to accumulate in the soma of spinal motor neurons.^28^ Ongoing and future studies will extend our findings and assess alterations in the distribution and trafficking of mitochondria.

Besides alterations in mitochondrial trafficking, mitochondrial dysfunction has been widely described in spinal cord from ALS patients^26–28^ as well as in animal and cellular models of ALS.^22–25^ Our results here build on these findings and demonstrate that alterations in tau phosphorylation may underlie mitochondrial dysfunction in ALS, given that ALS SNs enriched in pTau-S396 increased oxidative stress, reduced mitochondrial length and volume, and altered mitochondrial connectivity *in vitro*. Consistent with these *in vitro* findings we also found a significant decrease in mitochondrial density and length in post-mortem ALS mCTX, suggesting an increase in mitochondrial fragmentation as reported previously.^22–23^ Our findings of decreased mitochondrial density and length in ALS post-mortem mCTX together with an increase in synaptic DRP1 levels further supports the notion of increased mitochondrial fission in ALS. Hyperphosphorylated tau has also been shown to impair mitochondrial dynamics through an abnormal interaction with the pro-mitochondrial fission GTPase DRP1.^20–21^ Therefore, the concomitant increase in synaptic pTau-S396 and DRP1 occurring across ALS subtypes further suggests that these alterations may be a common pathogenetic mechanism in ALS. Lastly, the involvement of hyperphosphorylated tau in inducing mitochondrial fission in ALS was further confirmed by our *in vitro* studies, wherein we demonstrated alterations in mitochondrial length, volume and connectivity following treatment with ALS SNs, enriched in both pTau-S396 and DRP1.

Studies utilizing Drp1 heterozygous mice (Drp1^+/-^) with impartial DRP1 depletion or treated pharmacologically with mdivi-1, a DRP1 inhibitor, demonstrate that DRP1 inhibition can prevent mitochondrial dysfunction and synapse loss in AD.^21, 41^ Similarly, we reported that silencing *DRP1* prevented ALS SNs-induced decreases in mitochondrial length and volume in SH-SY5Y cells. Importantly, we recapitulated the same mitochondrial morphologic protection by reducing tau levels with the specific tau degrader QC-01-175, which is able to degrade disease-relevant forms of tau and p-Tau, including pTau-S396, in neuronal cells derived from FTD patients.^34^ Here, we further confirm the therapeutic potential of QC-01-175, as we reported a significant decrease in pTau-S396 levels following QC-01-175 treatment in SH-SY5Y cells treated with ALS SNs. Additionally, we demonstrated that reducing pTau-S396 with QC-01-175 mitigates mitochondrial fragmentations and preventes increases in ROS levels induced by ALS SNs, thus further supporting the involvement of tau in mitochondrial dysfunction in ALS and, importantly, identifying a novel potential therapeutic target for ALS.

Targeting mitochondrial proteins, such as DRP1, is a great challenge given that mitochondria are involved in several vital cellular functions thus increasing the possibility of deleterious side-effects such as the ones reported following treatment with the putative DRP1 inhibitor mdivi-1.^42–43^ Furthermore, the clinical application of siRNA is limited by both their poor pharmacokinetic properties and the possible induction of off-target effects.^44^ Moreover, the Food and Drug Administration (FDA) has only approved two RNA interference (RNAi)-based therapies in the last 20 years.^45–46^ Therefore, targeting tau to improve the status of mitochondria may represent a more viable therapeutic strategy. Our findings suggest that reducing pTau-S396 levels lead to an improvement in mitochondrial dynamics and, in turn, mitochondrial function, providing a new approach to mitigate mitochondrial dysfunction. Given that tau mainly localizes at the neuronal level,^47^ targeting tau to improve the balance in mitochondrial dynamics may be associated with less deleterious off-target effects. Future studies are planned to further investigate the neuroprotective effects of QC-01-175 in animal models of ALS to determine its therapeutic potential.

Collectively, our results in a large cohort of human post-mortem mCTX suggest that hyperphosphorylated tau at S396 may underlie mitochondrial fragmentation in ALS by interacting with the pro-fission GTPase DRP1. Lastly, our data provide the groundwork to assess QC-01-175 as a novel potential therapeutic strategy to improve mitochondrial morphology and function, and, in turn, motor neuron survival in ALS.

## Material and Methods

All methods were carried out in accordance with the guidelines and regulations of Massachusetts General Hospital and approved by the Massachusetts General Hospital licensing committees.

### Human tissue samples

Post-mortem mCTX from control and ALS patient brains were provided by the Massachusetts Alzheimer’s Disease Research Center (ADRC) with approval from the Massachusetts General Hospital IRB (1999p009556) and by the University of Edinburgh MRC/Sudden Death Brain Bank (under ethical approvals 15-HV-016; 11/ES/0022). In total we assessed 47 ALS and 25 non-neurological control motor cortices. The mean age was 68.2 years (SD=15.7) for control subjects and 62 years (SD=10.9) for ALS subjects. The samples consisted of 56% males in the control group and 53.2% males in the ALS group. There were 20 limb onset and 14 bulbar onset ALS samples, and 3 were diagnosed with ALS/FTD. Five of the ALS cases were positive for *C9ORF72* repeat expansion, while a single case was positive for *SOD1* mutation. The genetic status of the other ALS samples was unknown. Lastly, 24 of the ALS cases demonstrated TDP-43 proteinopathy. Post-mortem interval (PMI) range was 2-104h for control and 4-102h for ALS (Suppl. Table 1).

### Synaptoneurosomes (SNs) fractionation

SNs fractionation was performed as previously reported^32^ with a few modifications. Human post-mortem mCTX (200 to 300mg) were homogenized in 500μL ice-cold Buffer A composed of 25mM Hepes pH 7.5, 120mM NaCl, 5mM KCl, 1mM MgCl2, 2mM CaCl2, 2mM dithiothreitol (DTT), 1mM NaF supplemented with phosphate and protease inhibitors cocktail. The homogenates were then passed through two layers of 80μm nylon filters (Millipore, MA) to remove tissue debris. Seventy μL aliquot was saved, mixed with 70μL H2O and 23μL 10% sodium dodecyl sulphate (SDS), passed through a 27-gauge needle to shear DNA, boiled for 5min, and centrifuged at 15,000g for 15min to prepare the total extract. The remainder of the homogenates was passed through a 5μm Supor membrane Filter (Pall Corp, Port Washington, NY) to remove organelles and nuclei, centrifuged at 1000g for 5min and both pellet and supernatant were saved. The supernatants were collected in small crystal centrifuge tubes and centrifuged at 100,000g for 45min at 4^ο^C to obtain the cytosolic fractions. The pellets were resuspended in Buffer B composed of 50mM Tris pH 7.5, 1.5% SDS and 2mM DTT, boiled for 5min, and centrifuged at 15,000g for 15min to obtain SNs. Bradford assay was used to determine protein concentration in total extract, cytosolic fraction, and SNs.

### Western blotting

Western blots were performed using previously described protocols.^48^ Briefly, 50μg of proteins was resuspended in sample buffer and fractionated on either a 4-12% bis-tris gel or a 10-20% tricine gel for 90min at 120V. Then, proteins were transferred to a PVDF membrane in an iBlot Dry Blotting System (Invitrogen, Thermo Fisher, MA), and the membrane was blocked with 5% milk in tris-buffered saline with Tween 20 (TBST) before immunodetection with the following primary antibodies: tau (1:1000, DAKO, Denmark), pTau-S396 (1:500, Abcam, MA), DRP1 (1:500, BD Bioscience, CA), pDRP1-S616 (1:500, Cell Signaling, MA), Mfn1 (1:500, Proteintech, IL), Mfn2 (1:500; Abcam, MA), OPA1 (1:500, BD Bioscience, CA), Tomm20 (1:500; Novus Biological, CO), PSD-95 (1:500; Cell Signaling, MA), GADPH (1:1000, Millipore Sigma, Termecula, CA), and β-Actin (1:1000; Cell Signaling, MA), overnight at 4°C. Primary antibody incubation was followed by washes and incubation with secondary antibody for 1h (HRP-conjugated goat anti-rabbit IgG, and HRP-conjugated goat anti-mouse; Jackson ImmunoResearch Laboratories, West Grove, PA). Protein bands were visualized using the ECL detection system (Thermo Fisher Scientific, MA).

### Cell culture

Human neuroblastoma SH-SY5Y cells were grown in Dulbecco’s Modified Eagle Medium (DMEM)/Nutrient Mixture F-12 (1:1) supplemented with 10% inactivated fetal bovine serum (FBS), 2mM L-glutamine, and 50μg/ml streptomycin and 50IU/ml penicillin. The cells were kept at 5% CO2 at 37°C. Before each experiment, SH-SY5Y cells were treated with 10ng/mL human recombinant tau-2N4R (tau), and 10ng/mL SNs from control and ALS human post-mortem mCTX for 24h.

### Human caspase-3 Enzyme-Linked Immunosorbent Assay (ELISA)

Cell viability was assessed using a Human caspase-3 (active) ELISA kit (#KHO1091; Thermo Fisher Scientific, MA) following manufacture’s protocols. Briefly, SH-SY5Y cells were treated with tau and SNs from control (n=5) and ALS mCTX (n=9) (10ng/mL/24h) and homogenized in the Cell Extraction Buffer provided by the kit. Samples were incubated with a caspase-3 detection antibody for 1h at RT, washed (4x), and incubated with a stabilized chromogen for 30min at RT in the dark. The reaction was stopped with a stop solution before absorbance was read at 450nm. Caspase-3 levels were calculated based on a standard curve after subtracting background absorbance.

### Aconitase activity assay

Aconitase activity was measured using a commercially available kit (#MAK051; Sigma-Aldrich, MO) following manufacturer’s protocols. Briefly, SH-SY5Y cells were treated with tau and SNs from control (n=4) and ALS mCTX (n=4) (10ng/mL/24h) and homogenized in ice-cold Assay Buffer. Next, samples were centrifuged at 800g for 10min at 4°C to remove insoluble material, and the supernatant was used to assess c-aconitase activity. To assess m-aconitase activity the supernatant was centrifuged at 20,000g for 15min at 4°C. The pellets were collected, dissolved in Assay Buffer, and sonicated for 20sec. After preparation, the samples were activated with the Aconitase Activation Solution on ice for 1h and then incubated with the Reaction Mix for 1h at 25°C in the dark. Lastly, the samples were incubated with the Developer Solution at 25°C for 10min and the absorbance was measured at 450nm. C- and m-aconitase activity was calculated following the kit’s instruction after subtracting background.

### Immunostaining and Image Analysis

SH-SY5Y cells plated on Millicell EZ SLIDE glass slides (Millipore Sigma, PEZG S0816) were treated with tau, and SNs derived from control (n=3) and ALS mCTX (n=3) (10ng/mL/24h). At the end of the treatment, cells were fixed with 4% paraformaldehyde for 20min prior to immunostaining. Cells were permeabilized with 0.15% Triton-X in PBS for 20 min, washed in PBS (5x), and then blocked in 7.5% BSA in PBS-T for 45min. Cells were then incubated for 90min in Tomm20 antibody (1:1000; #NBP2-67501, Novus Biologicals, CO), followed by 5 washes in PBS. Cells were then incubated for 1h in goat-anti-rabbit AlexaFluor 488 secondary antibody (1:200; #A11034, ThermoFisher Scientific, MA), followed by 5x PBS washes. Cells were then incubated for 30min in HCS CellMask Deep Red (2μg/mL; #H32721, ThermoFisher Scientific, MA), washed in PBS (2x), followed by 10min incubation in Hoechst 33342 (1μg/mL; # H3570, ThermoFisher Scientific, MA). Slides were then washed in PBS (3x), and coverslipped with Vectashield Antifade Mounting Medium (Vector Labs, H-1000) prior to imaging. Z-stacks were acquired on a Nikon A1SiR confocal microscope with 60x magnification and 0.3μm step size. At least 9 random images were captured for each condition with a Nikon A1R inverted confocal microscope using a 60x objective. Images were captured using DAPI (Hoechst), FITC (Tomm20), and Cy5 (HCS CellMask Deep Red) filter sets. Automatic 3D Deconvolution was then performed within NIS-Elements software. Background subtraction of generated .tiff files were performed in ImageJ (National Institutes of Health; Bethesda, MD). Automatic thresholding numbers to be used in subsequent reconstruction were determined in ImageJ for each channel: Hoechst = Default; Tomm20 = Default; HCS CellMask Deep Red = Renyi. These background-subtracted files were then uploaded into Imaris (Bitplane, Zurich, Switzerland) to create volumetric reconstructions of each individual channel. Surface reconstructions of each channel were generated using the Surface tool, with threshold values as determined by ImageJ previously. Mitochondrial length was determined using the BoundingBoxOO Length C settings, measuring the length of the longest principal axis of each individual mitochondrion. Mitochondrial volume of each individual mitochondria was also determined from the surface volumetric reconstructions. Length or volume of every mitochondria for each condition were input into GraphPad Prism 9.0 software to generate cumulative frequency plots. The Data Analysis Histogram function on Microsoft Excel was used to generate binned frequency plots of mitochondrial length and volume.

### Electron microscopy

mCTX blocks for electron microscopy were made as previously described.^49^ Tissue samples were obtained from motor cortex at autopsy, cut into slices of no more than 1mm thick and 1 mm wide containing all 6 layers of cortex, and fixed for 48h in 4% paraformaldehyde, 2.5% glutaraldehyde in phosphate buffer (0.1M). Samples were post-fixed in osmium tetroxide (1%) for 30 minutes, stained with uranyl acetate (1% in 70% ethanol), dehydrated, and embedded in Durcupan resin. Utlrathin sections were cut on a Reichert-Jung Ultracut E. Lastly, sections were imaged on a JEM1011 electron microscope (JEOL) and images were acquired with a digital camera from Advanced Microscopy Techniques, Inc (Danvers, MA) and processed using AMT V601 software. For mitochondria analysis, an average of 15 images were taken/case at 25000x magnification in a systemic, random fashion from control (n=4) and ALS (n=5) mCTX (Supp. Table 2). Mitochondria were identified by the presence of internal cristae and defined outer membrane. Mitochondrial density was calculated from generated .tiff files in ImageJ as average of at least eight images/case. Mitochondrial density for each condition was input into GraphPad Prism 9.0 software to generate individual value scatter plots. Mitochondrial lengths were calculated from generated .tiff files in ImageJ and length of individual mitochondria for each condition were input into GraphPad Prism 9.0 software to generate cumulative frequency plots. All mitochondria within at least six images per subject were measured to obtain mitochondrial length. All individual mitochondrial lengths were used to generate cumulative frequency plots, whereas mitochondrial lengths from every case image were averaged for binned analyses. The Data Analysis Histogram function on Microsoft Excel was used to generate binned frequency plots of mitochondrial length and volume.

### Co-immunoprecipitation (co-IP)

The co-IP experiments were performed as previously reported.^50^ Briefly, 75μg of proteins from control and ALS post-mortem mCTX homogenates was incubated with 5μL anti-DRP1 (BD Bioscience, CA), and GAL4 immunoprecipitation buffer, composed of 250mM NaCl, 5mM ethylenediaminetetraacetic acid (EDTA), 1% Nonidet P-40, 50mM Tris pH 7.5. After incubation at 4°C for 3.5h, magnetic protein A beads (Invitrogen, Thermo Fisher, MA) were added (20μL per sample) and left in agitation overnight at 4°C. Next, samples were placed on a magnetic rack to remove the supernatants and washed in GAL4 buffer (3x). Twenty μL sample buffer was added in each sample before boiling at 95°C for 5min. For the input samples, used as control, 40μg of proteins from the same mCTX homogenates was resuspended in sample buffer and boiled at 95°C for 5min. The samples were then loaded in a 4-20% glycine gel as above described for the western blot method. An antibody against pTau-S396 (1:500; Abcam, MA) was used to immunoblot.

### Small interfering RNA (siRNA)

DRP1 knock down was obtained with siGENOME Human DNM1L siRNA (siDRP1) (SMARTPool; M-012092-01-0005; Horizon Discovery, Waterbeach, UK), using a siGENOME non-targeting control pool (SMARTPool; D-001206-13-05) as control. Transfection of SH-SY5Y cells was carried out in Optimem medium (ThermoFisher Scientific, MA) using Lipofectamine 2000 (ThermoFisher Scientific, MA) together with 10, 100, or 150nM siDRP1 for 24 and 48h. Western blot experiments were performed to verify the efficiency of the transfection by assessing DRP1 levels.

### QC-01-175 treatment

QC-01-175 was designed using a targeted protein degradation technology to recognize both tau and Cereblon (CRBN), a substrate receptor for the E3-ubiquitin ligase in order to specifically induce pathogenic tau ubiquitination and degradation.^34^ SH-SY5Y cells were treated with 1 or 10μM QC-01-175 for 2, 4, and 24h following treatment with ALS SNs (10ng/mL/24h). Western blot experiments were performed to verify the efficiency of QC-01-175 by assessing pTau-S396 levels.

### Treatment of SH-SY5Y cells

*siDRP1.* SH-SY5Y cells were transfected with siControl (150nM) and siDRP1 (150nM) for 24h, as previously described, and then treated with SNs from ALS mCTX (n=4) (10ng/mL) for 24h. At the end of the treatments, DRP1 levels were assessed western blots. Mitochondrial morphological parameters were measured as above-reported in SH-SY5Y cells treated with either recombinant tau protein (10ng/mL) or SNs derived from control (n=3) and ALS mCTX (n=3) for 24h following transfection with siControl (150nM) and siDRP1 (150nM) for 24h.

*QC-01-175.* SH-SY5Y cells were treated with ALS SNs (n=4) (10ng/mL) for 24h followed by exposure to QC-01-175 (10μM) for 4h. At the end of the treatments, pTau-S396 levels were assessed by western blots. Mitochondrial morphological parameters were measured as previously described in SH-SY5Y cells treated with either recombinant tau protein (10ng/mL) or SNs derived from control (n=3) and ALS mCTX (n=3) for 24h followed by exposure to QC-01-175 (10μM) for 4h. Reactive oxygen species were measured in SH-SY5Y cells treated with either tau, control (n=4) or ALS SNs (n=9) (10ng/mL/24h) followed by treatment with QC-01-175 (10μM/4h).

### Oxidative stress detection

Reactive oxygen levels were measured in SH-SY5Y cells using CellROX Orange Reagent, a fluorogenic probe for measuring oxidative stress in live cells (#C10443; Thermo Fisher Scientific, MA). Briefly, following specific treatments, the cells were incubated with 2.5μM CellROX for 30min at 37°C after which culture media was replaced for imaging on a BioTek Cytation 5 imaging reader (BioTek, VT). Images were captured from 6 random areas of each well every 10min for 2h. To analyze data generated by Cytation, images were processed using a custom Fiji macro, which opened each image and automatically calculated mean fluorescent intensity (MFI). The MFI values were then grouped using a custom Python script. Slopes (MFI/min) of CellRox fluorescence were determined and normalized to the slope of vehicle-treated cells for each run. Statistics and plots were then created using Graphpad Prism 9.0. All code for processing is currently hosted and publicly available at https://github.com/bendevlin18/cytation-analysis-2020.

### Statistics

Normal distributions of data were not assumed regardless of sample size or variance. Individual value plots with the central line representing the median and the whiskers representing the interquartile range, box plots with the central line representing the median, the edges representing the interquartile ranges, and the whiskers representing the minimum and maximum values, and scatter plots with the central line representing the mean, and the whiskers representing the standard deviation (SD) were used for graphical representation. Comparisons were performed using a non-parametric Mann-Whitney U test, a one-way ANOVA followed by Tukey’s post-hoc test, and a two-way ANOVA followed by Tukey’s post-hoc test at a significance level (α) of 0.05. Cumulative frequency distribution graphs were compared by Kolmogorov-Smirnov test (electron microscopy) or Kruskal-Wallis test (confocal microscopy). Exact p values (two-tailed) are reported.

### Study approval

The study was approved by the Mass General Brigham Healthcare Institutional Review Board (IRB). Written informed consent was obtained from all participants prior to study enrollment. Post-mortem consent was obtained from the appropriate representative (next of kin or health care proxy) prior to autopsy.

## Acknowledgments

T.P. was supported by an award from the Judith and Jean Pape Adams Charitable Foundation and Byrne Family Endowed Fellowship in ALS Research. S.M.K.F. was supported by the ALS Canada Tim E. Noël Postdoctoral Fellowship. S.D. was supported by the Alzheimer’s association (2018-AARF-591935) and the Jack Satter Foundation. D.H.O. is a recipient of an Alzheimer’s Association Clinician Scientist Fellowship (2018-AASCF-592307) and a Jack Satter Foundation Award; he is partially supported by the Dr. and Mrs. E. P. Richardson, Jr Fund for Neuropathology at MGH. S.J.H. was supported by the Alzheimer’s Association/Rainwater Foundation Tau Pipeline Enabling Program and the Stuart & Suzanne Steele MGH Research Scholars Program. The Massachusetts Alzheimer’s Disease Research Center is supported by the National Institute on Aging NIA (Grant P30AG062421). The Philly Dake Electron Microscopy Facility was supported by the Dake Family Foundation and by the NIH grant (1S10RR023594S10) to M.D. The authors would like to thank the patients and their families for sample donations.

## Author Contributions

T.P. and E.A.B. contributed to the study design, data collection, data analysis, and drafting of the manuscript. A.N.M., S.E.K., E.S., B.A.D., A.A.O., S.M.K.F., A.C.A., S.D., P.M.D. contributed to the data collection, data analysis, and editing of the manuscript. C.H., D.H.O., A.N., B.T.H., T.S.J., S.D.B., K.V., M.E.C., J.D.B., M.D., M.C.S., S.J.H. contributed to the study design, and editing of the manuscript. G.S.V. contributed to the study design, data analysis, drafting of the manuscript.

---

**Supplementary Figure 1.**
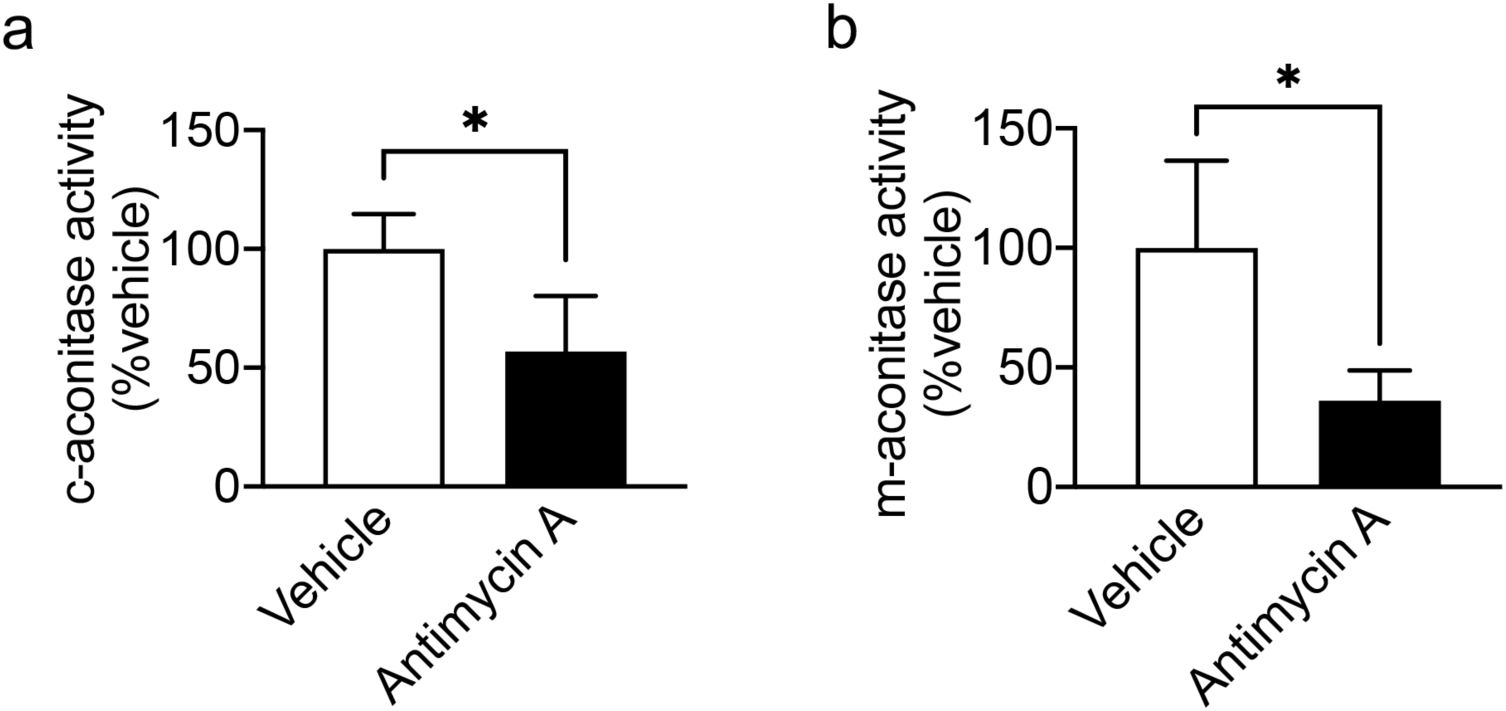
Antimycin A decrease c- and m-aconitase activity. (**a**) Antimycin A reduced (**a**) c-aconitase and (**b**) m-aconitase activity in SH-SY5Y cells (Mann-Whitney U test=0, p=0.0286, and Mann-Whitney U test=0, p=0.0286, respectively). *p<0.05.

**Supplementary Figure 2.**
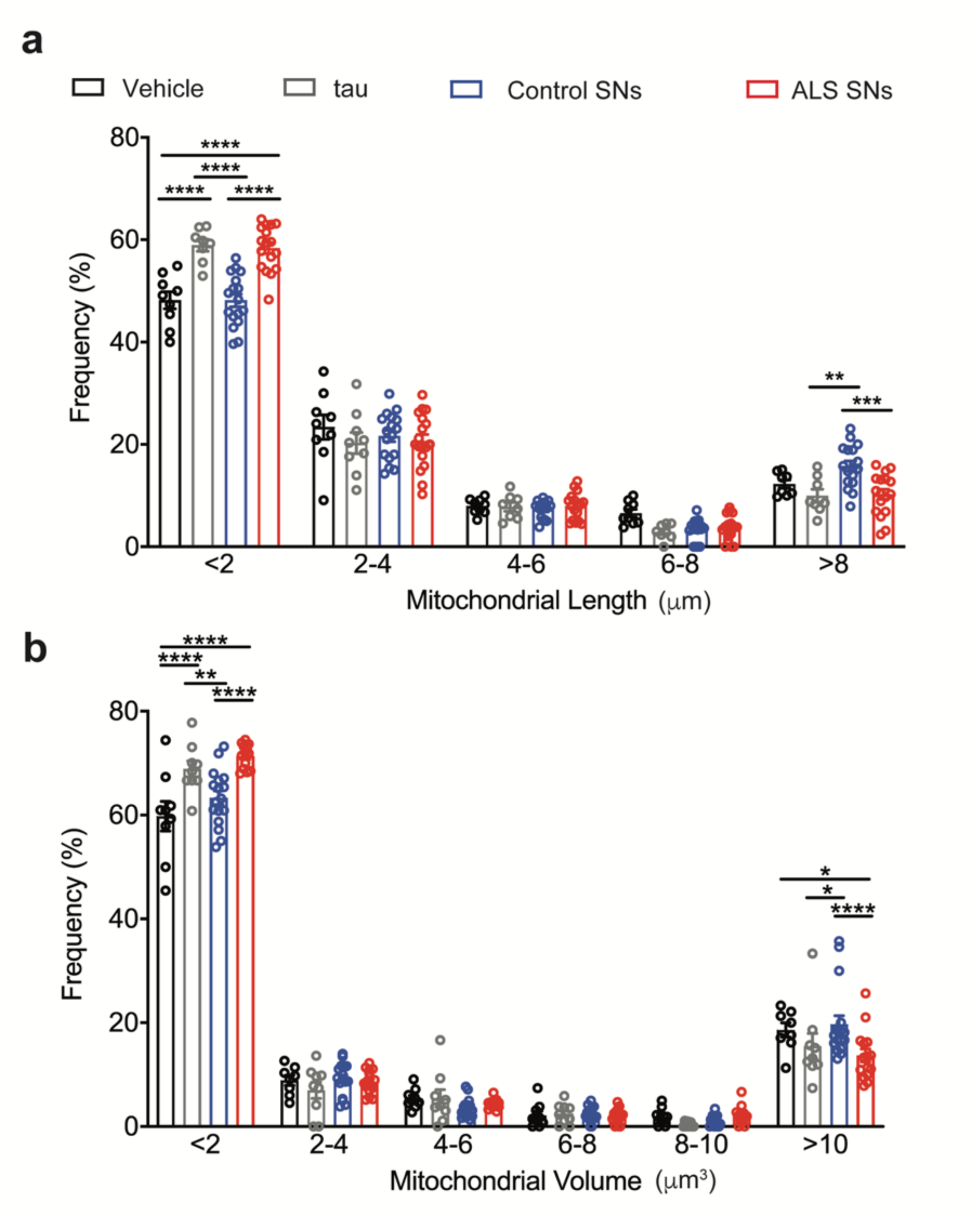
ALS SNs induce mitochondrial fragmentation. (**a**) Two-way ANOVA revealed an effect of length ([F(4,234)=1148], p<0.0001) and length X treatment interaction ([F(12,234)=9.629], p<0.0001) in SH-SY5Y cells. Tukey’s test revealed an increase in the frequency of smaller mitochondria (<2μm) in recombinant tau- and ALS SNs-treated cells compared to vehicle- (p<0.0001, and p<0.0001, respectively) and control SNs-treated cells (p<0.0001, and p<0.0001, respectively) as well as a decrease in the frequency of larger mitochondria (>8μm) in recombinant tau- and ALS SNs-treated cells compared to control SNs-treated cells (p=0.0041, and p=0.0003, respectively). (**b**) Two-way ANOVA demonstrated an effect of volume ([F(5,266)=1878], p<0.0001) and volume X treatment interaction ([F(15,266)=5.943], p<0.0001) in SH-SY5Y cells. Tukey’s test revealed an increase in the frequency of smaller mitochondria (<2μm^3^) in recombinant tau- and ALS SNs- compared to vehicle- (p<0.0001, and p<0.0001, respectively) and control SNs-treated cells (p=0.0028, and p<0.0001, respectively) as well as a decrease in the frequency of larger mitochondria (>10μm^3^) in ALS SNs- compared to vehicle-treated cells (p=0.0178) and in recombinant tau- and ALS SNs- compared to control SNs-treated cells (p=0.0418, and p<0.0001, respectively). *p<0.05; **p<0.01; ***p<0.001; ****p<0.0001

**Supplementary Figure 3.**
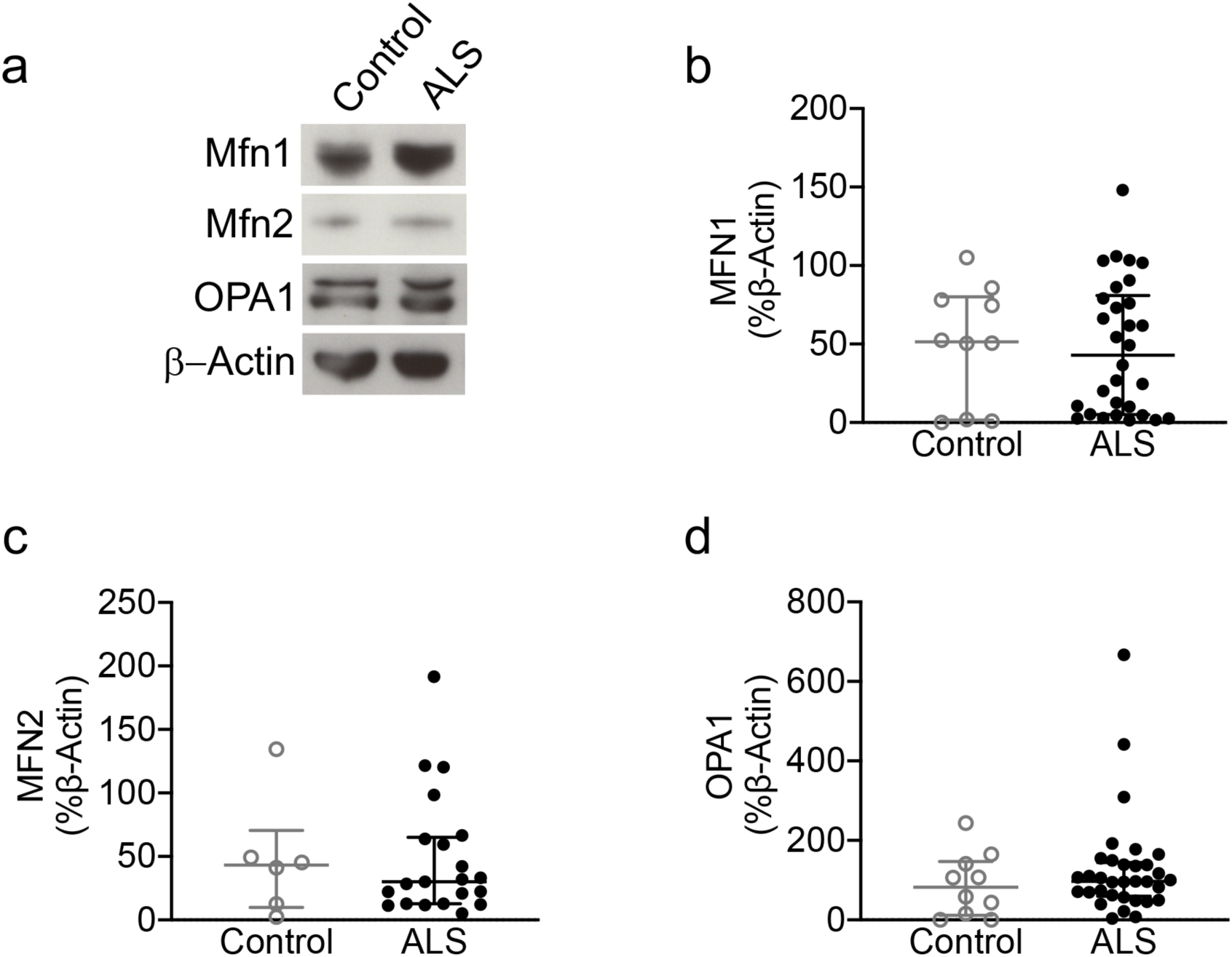
Pro-fusion proteins are not altered in ALS SNs. (**a**) Representative western blot images of Mfn1, Mfn2 and OPA1 in control and ALS SNs. (**b**) There was no significant change in Mfn1 (Mann-Whitney U test=144, p=0.8597) (**c**) Mfn2 (Mann-Whitney U test=61, p=0.9321) or (**d**) OPA1 levels (Mann-Whitney U test=135, p=0.3994) in ALS SNs (n=32) compared to controls (n=12).

**Supplementary Figure 4.**
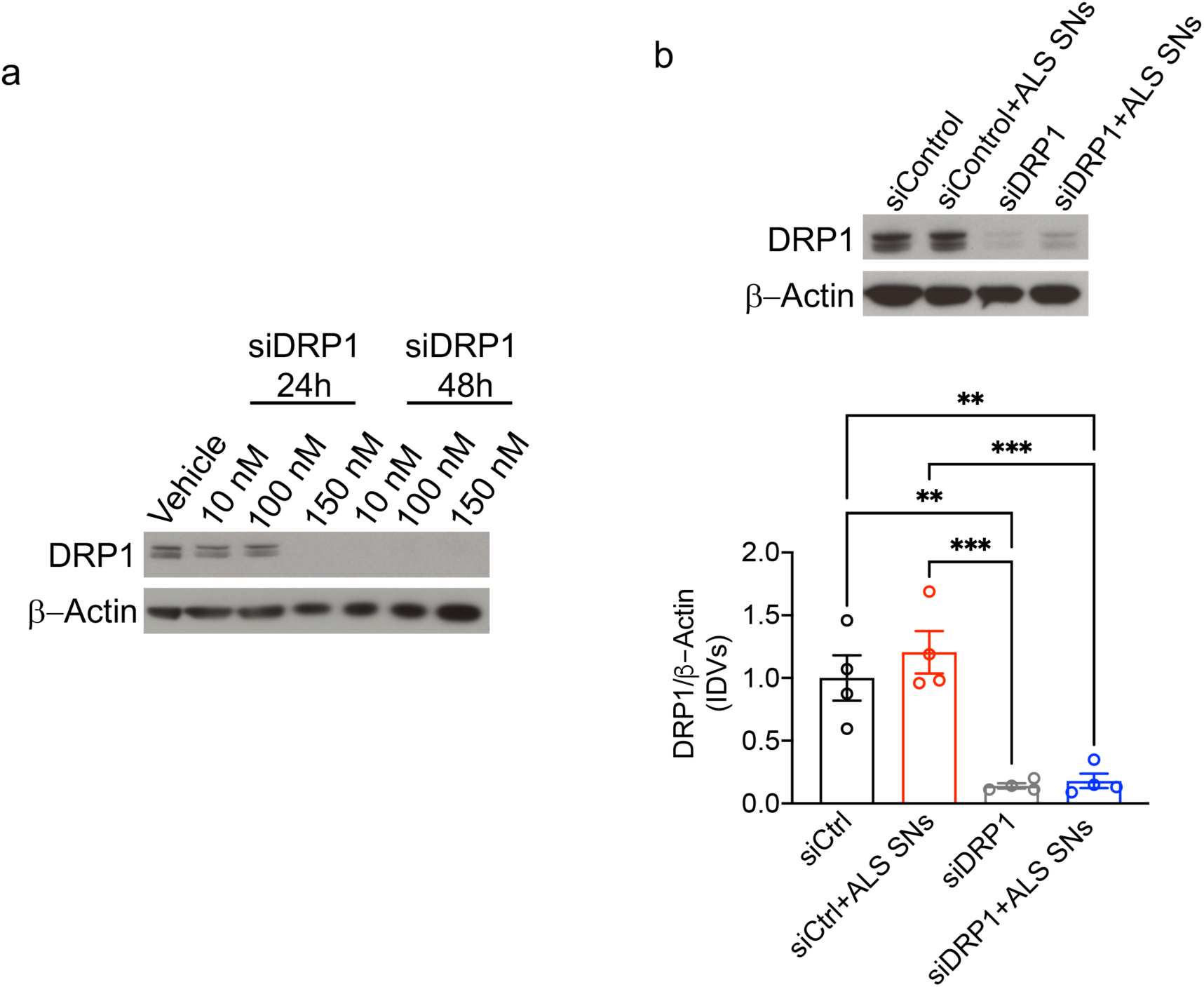
siDRP1 decreases DRP1 levels in ALS SNs-treated cells. (**a**) Representative western blot images of DRP1 in SH-SY5Y cells following siDRP1. DRP1 levels were reduced following transfection with 150nM siDRP1 for 24h and following 48h treatment. (**b**) There was a significant effect of treatment on DRP1 levels in SH-SY5Y cells (one-way ANOVA, [F(3,12)=18.52], p<0.0001) with a significant decrease in DRP1 levels in siDRP1- and siDRP1+ALS SNs-treated cells compared to siControl- (Tukey’s test, p=0.0023, and p=0.0033, respectively) and siControl+ALS SNs- treated cells (Tukey’s test, p=0.00004, and p=0.0005, respectively). **p<0.01; ***p<0.001.

**Supplementary Figure 5.**
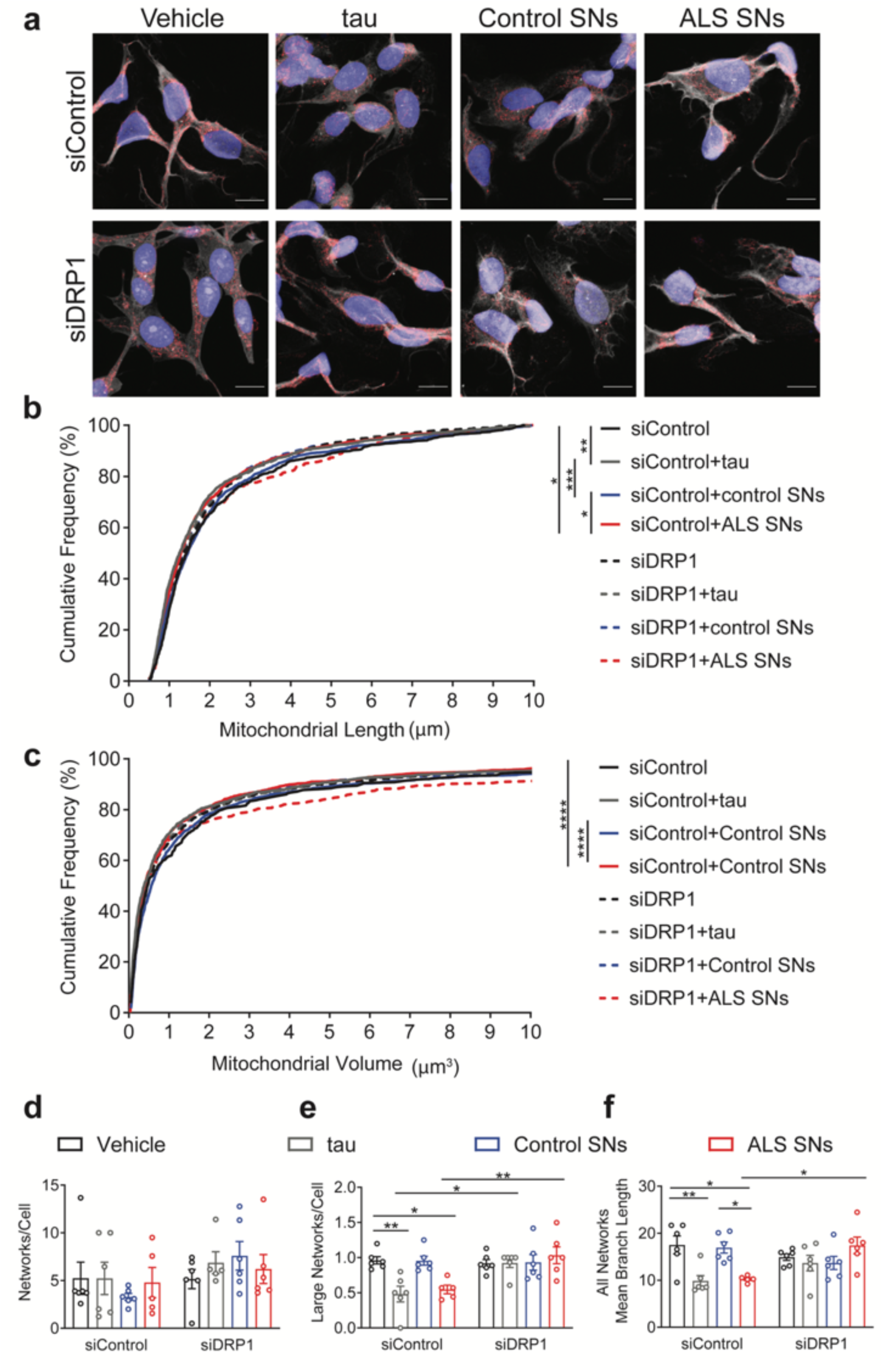
siDRP1 mitigates ALS SNs-induced alteration in mitochondrial connectivity. (**a**) Representative images of siControl- and siDRP1-transfected SH-SY5Y cells treated with vehicle, recombinant tau, control and ALS SNs stained with Hoechst (blue), CellMask (white), and Tomm20 (red). (**b**) Cumulative frequency graph indicated an effect of treatment on length (Kruskal-Wallis, H=33.14, p<0.0001) with smaller mitochondria in siControl+tau- and siControl+ALS SNs-treated (n=3) compared to siControl- (p=0.0023, and p=0.0477, respectively) and siControl+control SNs-treated (n=3) cells (p=0.0008, and p=0.0377, respectively). (**c**) Cumulative frequency graph indicated an effect of treatment on volume (Kruskal-Wallis, H=40.04, p<0.0001) with smaller mitochondria in siControl+tau- and siControl+ALS SNs- compared to siControl+control SNs-treated cells (p<0.00001, and p<0.0001, respectively). (**d**) There was no effect of treatment on networks/cell following treatments (two-way ANOVA, [F(3,38)=0.9047], p=0.4479). (**e**) Two-way ANOVA revealed an effect of treatment ([F(3,39)=3.953], p=0.0149), siDRP1 ([F(1,39)=12.80], p=0.0009), and treatment X siDRP1 interaction ([F(3,39)=5.659], p=0.0026) on large networks/cell in SH-SY5Y cells. Tukey’s test revealed a decrease in tau- and ALS SNs-compared to vehicle- (p=0.0056, and p=0.0374, respectively) or control SNs-treated cells (p=0.0064, and p=0.0416, respectively). siDRP1 prevented tau- and ALS SNs-induced decrease in large networks/cell (Tukey’s test, p=0.0137, and p=0.0082, respectively). (**f**) Two-way ANOVA demonstrated an effect of treatment ([F(3,39)=4.034], p=0.0136) and treatment X siDRP1 interaction ([F(3,39)=6.527], p=0.0011) on mean branch length. Tukey’s test demonstrated a decrease in siControl+tau and siControl+ALS SNs-compared to siControl- (p=0.0068, and p=0.0195, respectively) or siControl+control SNs-treated cells (p=0.0153, and p=0.0398, respectively). siDRP1 prevented ALS SNs-mediated branch length alterations (p=0.0398). *p<0.05; **p<0.01; ***p<0.001; ****p<0.0001.

**Supplementary Figure 6.**
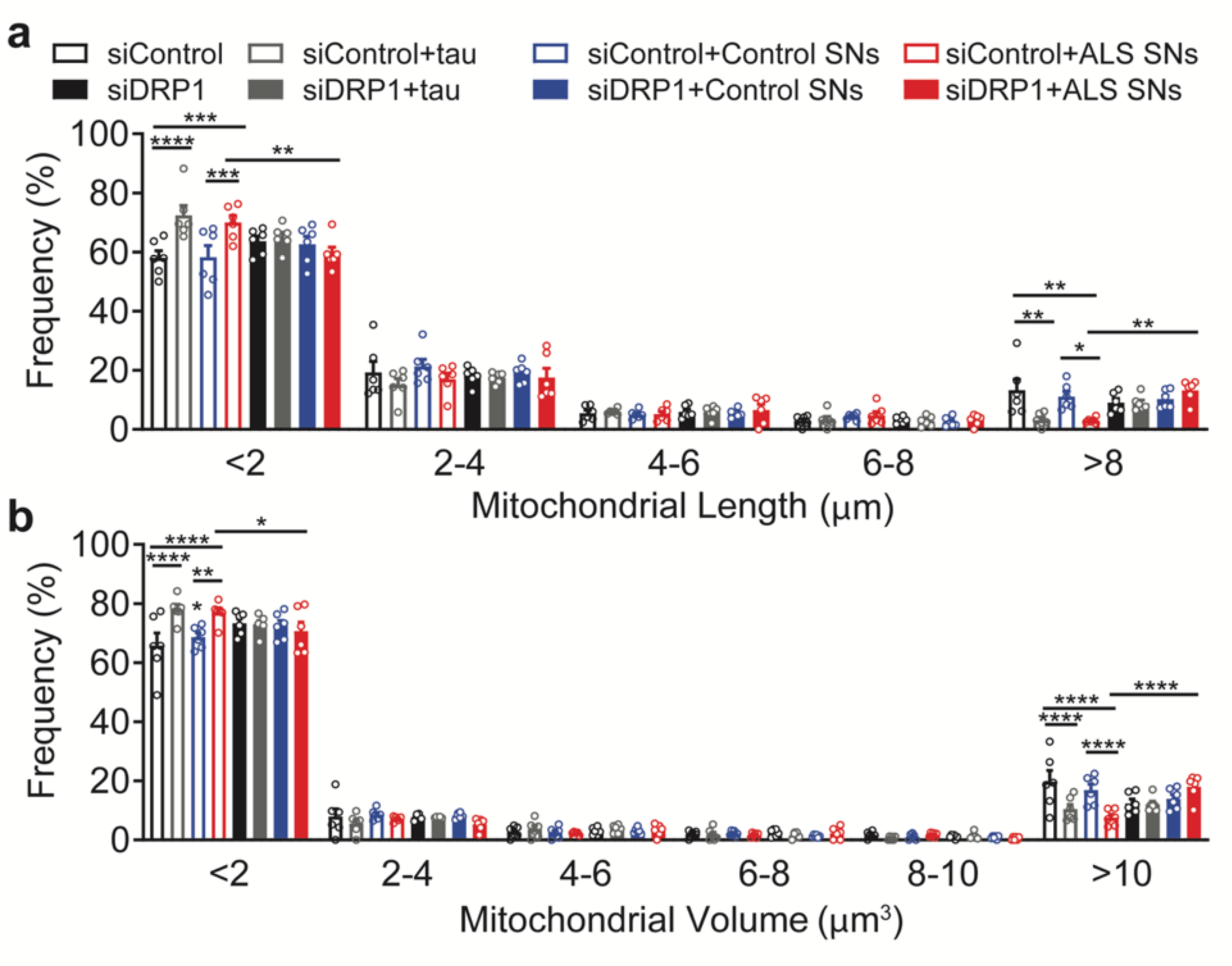
siDRP1 prevents ALS SNs-induced mitochondrial fragmentation. (**a**) There was a significant effect of length (two-way ANOVA, [F(4,200)=1502], p<0.0001) and length X treatment interaction (two-way ANOVA, [F(28,200)=3.621], p<0.0001) in SH-SY5Y cells. Tukey’s test revealed a significant increase in the frequency of smaller mitochondria (<2μm) in siControl+tau- and siControl+ALS SNs- (n=3) compared to siControl (p<0.0001, and p=0.0002, respectively) and siControl+control SNs-treated (n=3) cells (p<0.0001 and p=0.0002, respectively). Similarly, Tukey’s test demonstrated a significant decrease in the frequency of larger mitochondria (>8μm) in recombinant tau- and ALS SNs-compared to siControl-treated cells (p=0.0032 and p=0.0024, respectively) as well as in ALS SNs-compared to control SNs-treated cells (p=0.0433). siDRP1 prevented ALS SNs-induced increases in smaller mitochondria (p=0.0015) and reductions in larger mitochondria (p=0.0433). (**b**) There was a significant effect of both mitochondrial volume (two-way ANOVA, [F(5,237)=3239], p<0.0001) and volume X treatment interaction (two-way ANOVA, [F(35,237)=3.702], p<0.0001) in SH-SY5Y cells. Tukey’s test revealed a significant increase in the frequency of smaller mitochondria (<2μm^3^) in siControl+tau- and siControl+ALS SNs compared to siControl (p<0.0001, and p<0.0001, respectively) and control SNs-treated cells (p<0.0001 and p=0.0007, respectively). Similarly, Tukey’s test demonstrated a significant decrease in the frequency of larger mitochondria (>10μm^3^) in recombinant tau- and ALS SNs-treated cells compared to siControl (p<0.0001) as well as compared to siControl+control SNs-treated cells (p=0.0203 and p<0.0001, respectively). siDRP1 prevented ALS SNs-induced increases in smaller mitochondria (p=0.0289) and reductions in larger mitochondria (p<0.0001). *p<0.05; **p<0.01; ***p<0.001; ****p<0.0001.

**Supplementary Figure 7.**
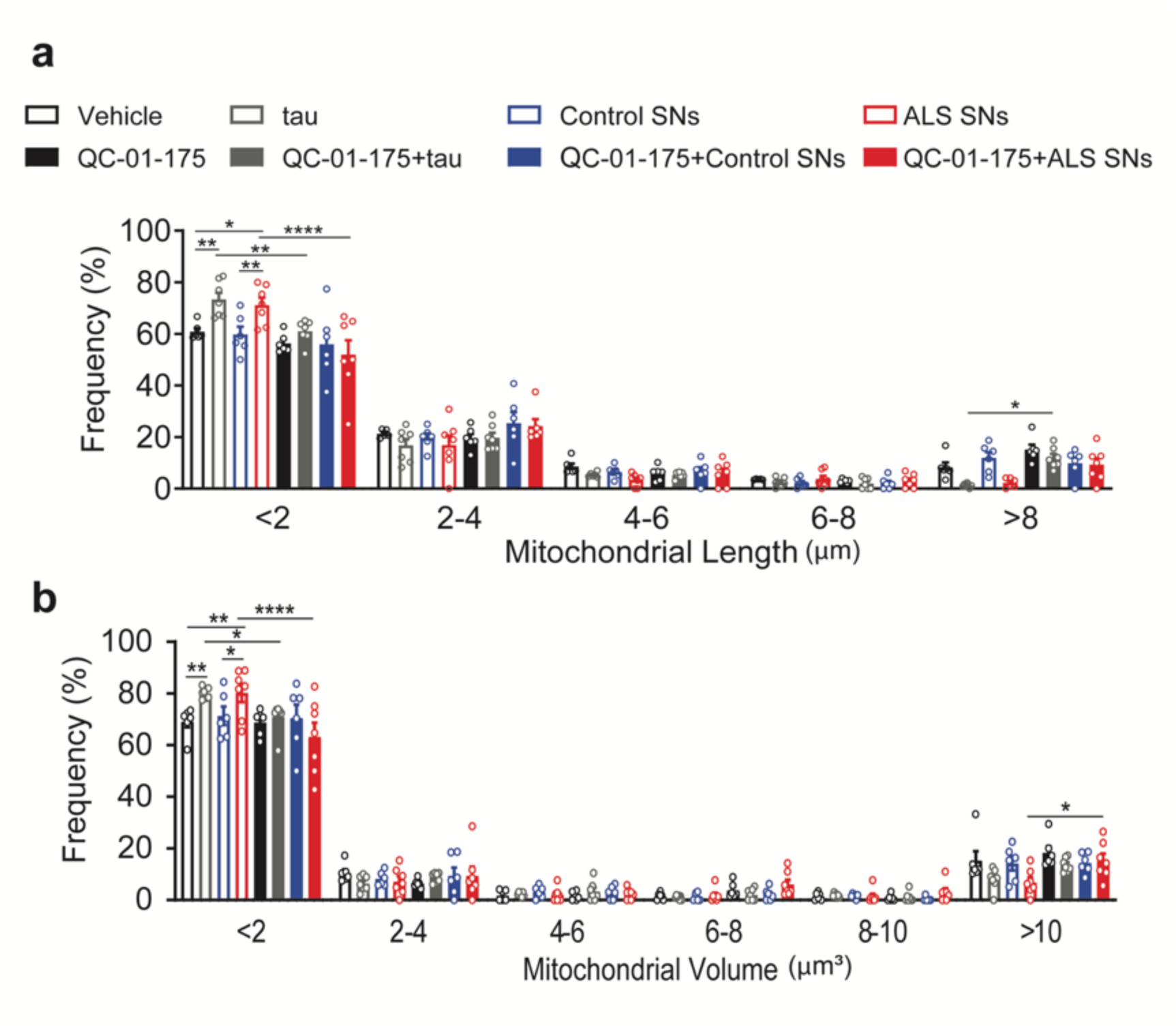
QC-01-175 prevents ALS SNs-induced mitochondrial fragmentation. (**a**) There was a significant effect of length (two-way ANOVA, [F(4,215)=955.7], p<0.0001), and length X treatment interaction (two-way ANOVA, [F(28,215)=4.756], p<0.0001). Tukey’s test revealed an increase in the frequency of smaller mitochondria (<2μm) in recombinant tau- or ALS SNs- (n=3) compared to vehicle- (p=0.0021, and 0.0235, respectively) and control SNs-treated (n=3) cells (p=0.0005, and p=0.0068, respectively). QC-01-175 prevented recombinant tau- and ALS SNs-induced increase in the frequency of smaller mitochondria (p=0.0014 and p<0.0001, respectively). (**b**) There was a significant effect of volume (two-way ANOVA, [F(5,264)=1510], p<0.0001), and volume X treatment interaction (two-way ANOVA, [F(35,264)=2.864], p<0.0001) in SH-SY5Y cells. Tukey’s test revealed an increase in the frequency of smaller mitochondria (<2μm^3^) in recombinant tau- and ALS SNs- compared to vehicle- (p=0.0034, p=0.0027, respectively) and control SNs-treated cells (p=0.0449, and p=0.0373, respectively). QC-01-175 prevented recombinant tau- and ALS SNs-induced increase in the frequency of smaller mitochondria (p=0.0179 and p<0.0001, respectively) as well as ALS SNs- induced decrease in the frequency of larger mitochondria (p=0.0470). *p<0.05; **p<0.01; ****p<0.0001.

**Supplementary Figure 8.**
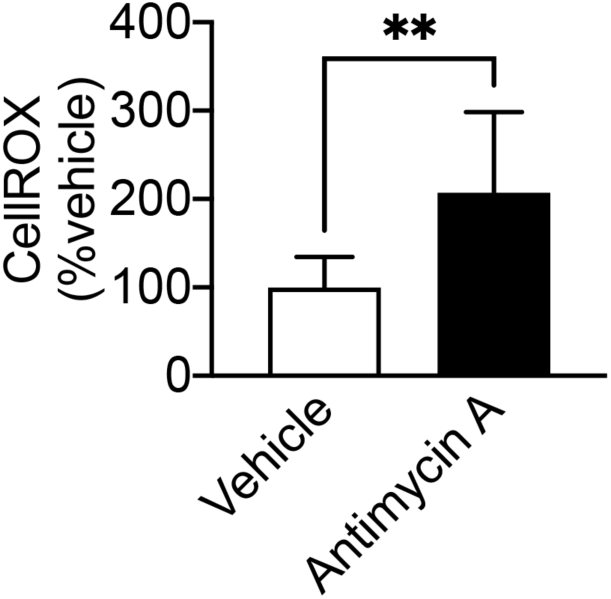
Antimycin A decreases ROS levels. Antimycin A decreased ROS levels in SH-SY5Y cells (Mann-Whitney U test=7, p=0.0037). **p<0.01.

**Supplementary Table 1.** **Human post-mortem mCTX information.**

|  | Sex |  | Onset site |  | Genotype |  | TDP-43<br>proteinopathy | FTD | Age at disease<br>onset<br>(average, SD) | Age of death<br>(average, SD) | PMI (h) |
| --- | --- | --- | --- | --- | --- | --- | --- | --- | --- | --- | --- |
|  | M | F | Bulbar | Limb | C9ORF72 | SOD1 |  |  |  |  |  |
| **Control** | 14 | 11 | N/A |  | N/A |  | N/A | N/A | N/A | 62.25 (15.3) | 2-104 |
| **ALS** | 26 | 21 | 14 | 20 | 5 | 1 | 24 | 3 | 60.84 (SD=11.6) | 62.51 (11.1) | 4-102 |

**Supplementary Table 2.**
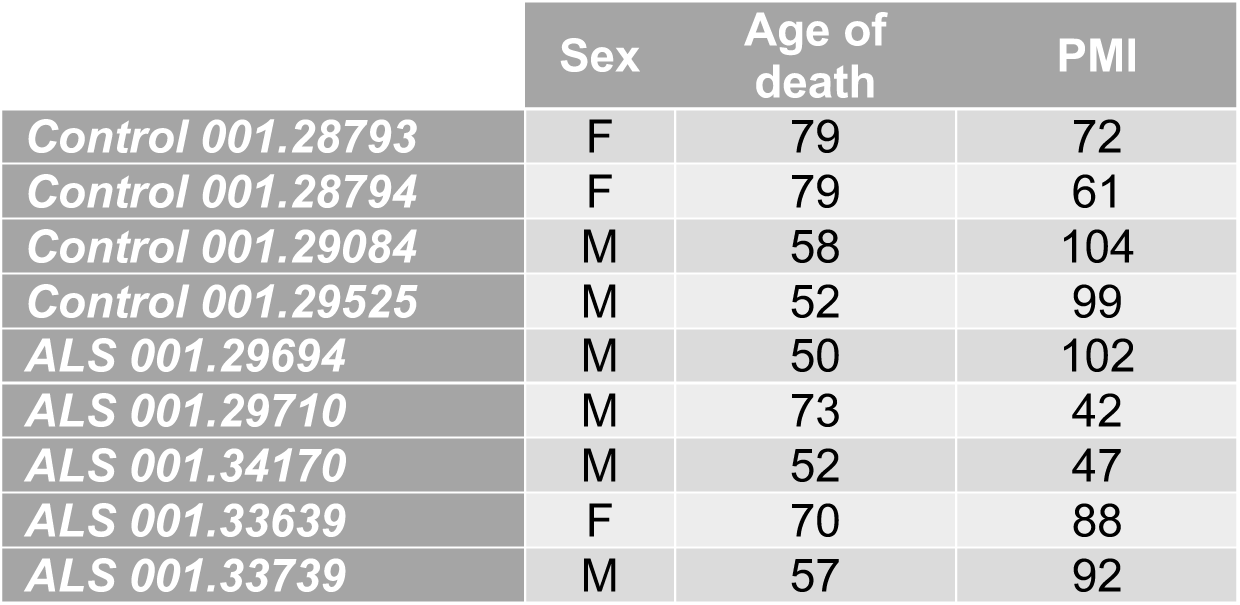
**EM blocks from human post-mortem mCTX.**

